# FTSH3 facilitates complex I degradation through a direct interaction with the complex I subunit PSST

**DOI:** 10.1101/2023.02.23.529725

**Authors:** Abi S. Ghifari, Aneta Ivanova, Oliver Berkowitz, James Whelan, Monika W. Murcha

**Affiliations:** School of Molecular Sciences & ARC Centre of Excellence in Plant Energy Biology, The University of Western Australia, 35 Stirling Highway, Crawley, Perth WA 6009, Australia; Department of Animal, Plant and Soil Science, School of Life Science, ARC Centre of Excellence in Plant Energy Biology, La Trobe University, Bundoora VIC 3086, Australia; College of Life Science, Zhejiang University, Hangzhou, Zhejiang 310058, P.R. China

## Abstract

Complex I (NADH dehydrogenase), the largest complex involved in mitochondrial oxidative phosphorylation is composed of nuclear and mitochondrial encoded subunits. Its assembly requires sequential addition of subdomains and modules. As it is prone to oxidative damage, complex I subunits continually undergo proteolysis and turnover. We describe the mechanism by which complex I abundance is regulated in a complex I deficient mutant. Using a forward genetic approach we have identified that the complex I Q-module domain subunit PSST, interacts with FTSH3 (Filamentous Temperature Sensitive H3) to mediate the disassembly of the matrix arm domain module for proteolysis and turnover as a means of protein quality control. We show the direct interaction of FTSH3 with PSST and identify the amino acid residues required for this interaction. It is the ATPase function of FTSH3 that is required for the interaction, as the mutation can be compensated by a proteolytically inactive form of FTSH3. Furthermore, it cannot be compensated by FTSH10 that is also located in mitochondria, as the latter does not interact with PSST. This study reveals the mechanistic process at the resolution of the residues involved of how FTSH3 recognises complex I for degradation.

## Introduction

Mitochondrial ATP is produced through oxidative phosphorylation (OXPHOS) by the combined action of multi-subunit protein complexes on the inner membrane and the mobile electron carriers ubiquinone and cytochrome *c*. Five distinct protein complexes are located in the mitochondrial inner membrane: NADH::ubiquinone oxidoreductase – complex I (CI), succinate dehydrogenase – complex II (CII), cytochrome *bc_1_* – complex III (CIII), and cytochrome *c* oxidase – complex IV (CIV), and ATP synthase. NADH generated from the tricarboxylic acid (TCA) cycle is oxidised by CI with subsequent transfer of electrons via mobile carriers through ubiquinone to CIII and cytochrome *c* to CIV with reduction of oxygen to water. This process creates a proton gradient necessary for ATP synthesis. CI is the first site of electron transfer and also the largest complex of OXPHOS pathway, consisting of 47-51 subunits in plants (Soufari et al., 2020; Klusch et al., 2021, 2022; Maldonado et al., 2022). Functionally, plant CI can be divided into four modules: NADH-binding module (N), ubiquinone-binding module (Q), proton-pumping (P) module, and carbonic anhydrase (CA) module. NQ modules form the hydrophilic matrix arm domain that protrudes into the matrix, which contains subunits involved in electron transfer from NADH to ubiquinone. The membrane arm is composed of the proximal (P_P_) and distal proton-pumping (P_D_) modules, and the CA module is matrix facing anchored to the inner membrane. As such CI assembly requires a sequential multistep assembly pathway, assisted by various assembly factors to form sub-modules and assembly intermediates (Schimmeyer et al., 2016; Ivanova et al., 2019; Ligas et al., 2019; Maldonado et al., 2020; Soufari et al., 2020).

The matrix arm domain is essential for overall CI activity containing essential cofactors. This includes a flavin mononucleotide (FMN) prosthetic group covalently bound in 51 kD subunit for NADH binding, an ubiquinone-binding domain formed by NAD1 and PSST, and eight iron-sulfur (Fe-S) clusters along NQ-modules required for electron transfer (Soufari et al., 2020). Redox reactions and electron transfer generates reactive oxygen species (ROS) (Hirst and Roessler, 2016), which renders the matrix arm more susceptible to oxidative damage (Szczepanowska et al., 2020). Consequently, matrix arm domain subunits have been shown to exhibit some of the highest protein turnover rates compared to the membrane module subunits, and compared to other mitochondrial proteins (Nelson et al., 2013; Li et al., 2017; Szczepanowska et al., 2020). The observed selective turnover of matrix arm domain subunits suggests that a submodule disassembly pathway occurs, allowing for selective proteolysis of damaged subunits and thus conservation of energy.

Previous studies have implicated a number of mitochondrial proteases involved *in organello* protein degradation. Plant mitochondria harbour several classes of proteases that belong to the AAA+ (ATPase-associated with various cellular activity) superfamily, which require ATP for activity (van Wijk, 2015; Opalińska and Jańska, 2018; Ghifari and Murcha, 2022). Mitochondrial AAA+ proteases such as LON1 (long filamentous phenotype-1), CLP (caseinolytic protease), and FTSH (filamentous temperature sensitive H) proteases have all been previously reported to be involved in OXPHOS complexes degradation and turnover (Li et al., 2017; Petereit et al., 2020; Huang et al., 2020; Ivanova et al., 2021). However, in contrast to the well-established cytosolic ubiquitin-proteasome degradation system (Vierstra, 2009), the mechanisms of OXPHOS complex disassembly, degradation, and turnover remains poorly understood.

Using a forward genetic screen in a CI defective mutant background *ciaf1* (complex I assembly factor 1) (Ivanova et al., 2019), we previously identified the inner membrane-bound AAA+ protease FTSH3 to play a role in regulating CI activity and abundance through the generation of a revertant named *rmb1 (restoration of mitochondrial biogenesis-1*) (Ivanova et al., 2021). This revertant displayed a restoration of CI activity and abundance, and biochemical and genetic analysis of *rmb1::ciaf1* determined that FTSH3 is involved in the disassembly of the matrix arm domain. Furthermore, it was shown that the ATPase domain of FTSH3, and not the proteolytic domain, was responsible for CI matrix arm disassembly in the CI defective background. The mutation in FTSH3 within the ATPase domain (P415L) was able to restore CI activity and abundance of matrix arm domain subunits. This restoration of phenotype was reverted by complementation with FTSH3. Interestingly, complementation was also achieved with a proteolytic inactive FTSH3 (FTSH3^TRAP^) indicating that the proteolytic function of FTSH3 was not responsible for CI abundance and activity and rather it was the ATPase function of FTSH3 that played a role with regards to CI (Ivanova et al., 2021).

However, the mechanism by which FTSH3 regulates CI disassembly was not elucidated in these studies. Here, we have determined that FTSH3 is involved in the disassembly of the matrix arm domain via a direct protein-protein interaction with PSST, a 20 kD subunit located at the interface of the membrane and matrix module (Soufari et al., 2020; Klusch et al., 2021, 2022; Maldonado et al., 2022). We identify and characterise an additional independent ethyl-methane-sulfonate (EMS) revertant of *ciaf1* that contains a mutation within the PSST gene (At5g11770) named *rmb2::ciaf1 (restoration of mitochondrial biogenesis-2). rmb2::ciaf1* displays a restoration of CI abundance, activity and consequently plant growth similarly to what we previously observed for the FTSH3 revertant mutant *rmb1::ciaf1* (Ivanova et al., 2021). Using a variety of protein-protein interaction assays we demonstrate a physical interaction between FTSH3 and PSST, that can be abolished by either *rmb1* and/or *rmb2* mutations (FTSH3^P415L^/PSST^S70F^). We observed an increased abundance of matrix arm subunits in *rmb2::ciaf1*, as we previously observed in *rmb1::ciaf1*, in addition to delayed protein turnover rates in both mutants. Therefore, combining genetic, biochemical, and proteomic analyses of two independent revertant lines that display restored CI abundance in a CI defective background, we have identified that FTSH3, via a direct interaction with the CI PSST subunit plays a role in the disassembly of the matrix arm domain of CI.

## Results

### The delayed growth phenotype in *ciaf1* was restored by a mutation in PSST

To identify regulators of CI biogenesis we carried out a forward genetic screen on a CI defective mutant. This mutant is a knockdown of the CI assembly factor-1 (CIAF1), a mitochondrial LYR domain-containing protein required for iron-sulfur (FeS) cluster insertion into matrix arm domain subunits (Ivanova et al., 2019). T-DNA insertional knockdowns of CIAF1 exhibit an almost complete lack of CI, due to the stalled assembly of the matrix arm domain and consequently, a severely delayed growth phenotype (Ivanova et al., 2019). This delayed growth phenotype, characteristic of many CI mutants, allowed for a visual screen of revertant mutants showing restored growth.

Here, we characterise an EMS mutant named *rmb2::ciaf1* that exhibits a substantial and significant growth restoration compared to *ciaf1* (**Figure 1A-B, Figure S1**). *rmb2::ciaf1 displays* significantly improved growth parameters, including plant height, number of rosette leaves, and maximum rosette radius compared to *ciaf1* throughout the growth cycle (**Figure 1A-B, Figure S1**). Whole genome sequencing of segregating homozygous *rmb2::ciaf1* populations identified the single nucleotide change of C to T at the chromosomal location of the gene designated At5g11770. The mutation corresponds to a single amino acid change from serine to phenylalanine at position 70 (S70F) in PSST, a 20 kD CI subunit that is located in the matrix arm Q-module in close proximity to the membrane arm submodule (Soufari et al., 2020; Klusch et al., 2021, 2022; Maldonado et al., 2022). PSST, also known as NADH dehydrogenase ubiquinone Fe-S protein-7 (NDUFS7) was shown to be involved in ubiquinone binding and reduction (Galemou Yoga et al., 2019; Soufari et al., 2020). The S70 residue is located at the N-terminal domain of the mature PSST protein, with the first 64 residues containing the N-terminal mitochondrial targeting peptide (mTP) (Soufari et al., 2020; Klusch et al., 2021) (**Figure 1C**).

**Figure 1.**
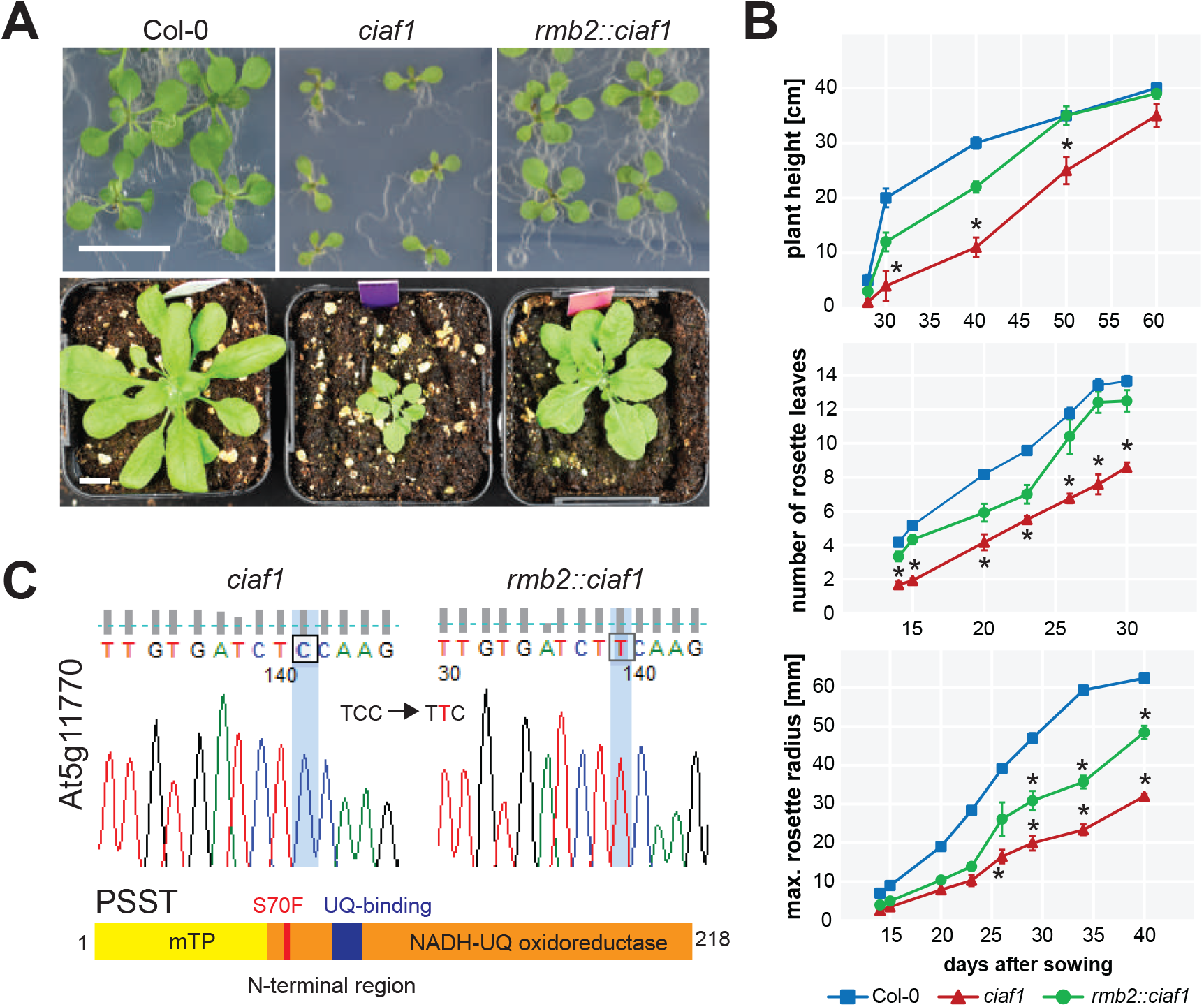
Identification of *rmb2::ciaf1*, an EMS-induced revertant *restoration of mitochondrial biogenesis-2 (rmb2*) in the COMPLEX I ASSEMBLY FACTOR-1 (CIAF1) knockdown *ciaf1* background. **A.** Phenotypic analysis of Columbia-0 (Col-0), *ciaf1* (SALK_143656) and *rmb2::ciaf1* on agar medium (half-strength Murashige-Skoog basal salt supplemented with 1.5% w/v sucrose) after 14 days (top panels) and on soil after 28 days (bottom panels), showing developmentally-delayed growth in *ciaf1* can be partially restored by mutation in *rmb2::ciaf1*. Scale bar = 1 cm. **B.** Growth parameter analyses: plant height, number of rosette leaves, and maximum rosette radius showing a restoration of growth and development in *rmb2::ciaf1* compared to *ciaf1*. Asterisks indicate statistically different datasets (mean ± SD, *n*=20, single-factor ANOVA *p*<0.05, followed by Tukey-Kramer post-hoc test). **C.** Whole genome sequencing of *rmb2::ciaf1* identifying a single-point mutation (TCC-TTC) within the At5g11770 gene, which corresponds to a mutation of position 70 serine to phenylalanine (S70F) in the N-terminal region of PSST, a 20 kD complex I Q-module domain subunit.

### Complex I activity and abundance of individual subunits is restored in *rmb2::ciaf1*

To confirm that the mutation in PSST is responsible for the observed phenotypic restoration, several crosses and complementation lines were generated (**Figure 2A)**. Firstly, the S70F mutation was introduced into a Col-0 background by filial crossing to produce *rmb2*::Col-0, resulting in no alterations to plant growth phenotype compared to Col-0 (**Figure 2A, Figure S1**). Secondly, a complementation line was generated by a stable transformation of *rmb2::ciaf1* with *PSST* under the control of its native promoter (*rmb2*^pro-PSST^) (**Figure 2A**). This complementation line exhibited in a reversion back to the small, developmentally delayed growth phenotype similar to *ciaf1* (**Figure 2A, Figure S1**), confirming that the PSST gene product is responsible for the observed phenotype in *rmb2::ciaf1*. Over-expression of *PSST* under the control of the cauliflower mosaic virus (CaMV) 35S promoter in Col-0 background was generated (*35S*::PSST), which displayed no significant alteration to plant growth parameters (**Figure 2A, Figure S1**).

**Figure 2.**
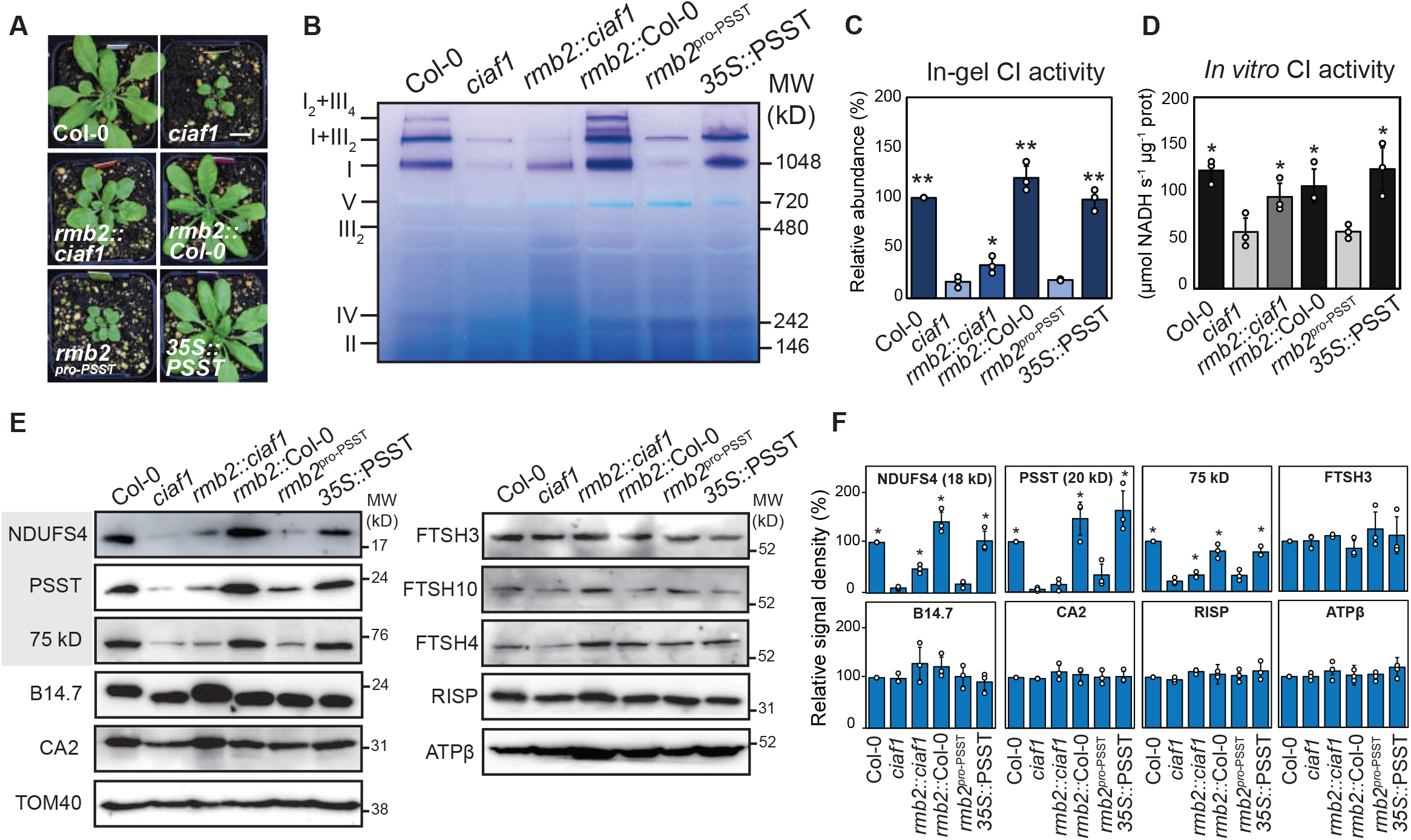
*rmb2::ciaf1* exhibits partially restored complex I activity and abundance. **A.** Phenotypic analysis of *rmb2::ciaf1, rmb2* in Col-0 background (*rmb2*::Col-0), *rmb2* complemented with PSST under a native promoter (*rmb2*::pro-PSST), and PSST overexpression line (*35S*::PSST). Scale bar = 1 cm. **B.** Blue-native PAGE followed by in-gel complex I activity staining of isolated mitochondria from all lines. The position and molecular weight of the OXPHOS complexes are indicated. **C.** Activity of complex I relative to Col-0 as quantified from in-gel staining in Figure 2B (mean ± SD, *n*=3 biological replicates). Statistically different datasets compared to *ciaf1* are indicated (* *p*<0.05, ** *p*<0.001, single-factor ANOVA). **D.** Assay of complex I activity measured *in vitro* from isolated mitochondria (mean ± SD, *n*=3 biological replicates). Statistically different datasets compared to *ciaf1* are indicated (* *p*<0.05, ** *p*<0.001, single-factor ANOVA) **E.** Immunodetection of isolated mitochondria from all lines, including CI NQ-module subunits (75 kD, NDUFS4, PSST, *highlighted gray*), other CI subunits (B14.7, CA2), other OXPHOS subunits (RISP, ATPβ), and mitochondrial AAA+ proteases (FTSH3, FTSH10, FTSH4). Outer membrane transport protein TOM40 was used as loading control. **F.** Densitometry measurements of immunoblot signals in Figure 2D relative to Col-0 for NDUFS4 (18 kD), PSST (20 kD), 75 kD, FTSH3, B14.7, CA2, RISP, and ATPβ normalised with TOM40 signal as loading control (mean ± SD, *n*=3 biological replicates). Statistically different datasets compared to *ciaf1* are indicated by asterisks * (single-factor ANOVA followed by Tukey-Kramer posthoc test).

Mitochondria were isolated from all genotypes and CI abundance was determined by blue-native polyacrylamide gel electrophoresis (BN-PAGE), followed by in-gel NADH dehydrogenase activity (Schertl and Braun, 2015). In-gel CI activity staining indicates a significant increase to CI abundance of ~2-fold in *rmb2::ciaf1* compared to *ciaf1* (**Figure 2B-C**) (*p*<0.05, *n*=3) whilst the complementation line (*rmb2*^pro-PSST^) resulted in a decrease of CI abundance back to the levels that are observed in *ciaf1* (**Figure 2B-C**). Introduction of *rmb2* in a Col-0 background (*rmb2*::Col-0) and PSST over-expression (*35S*::PSST) in a Col-0 background did not result in any changes to CI abundance compared to Col-0 (**Figure 2B-C**).

To confirm these results, an *in vitro* colorimetric plate-based assay was used to measure CI activity (Huang et al., 2015) revealing similar changes. A significant decrease of ~50% in CI activity was measured in *ciaf1* compared to Col-0 that was restored to ~80% of Col-0 *rmb2::ciaf1*. Complementation reverted activity back to to the levels observed in *ciaf1* (**Figure 2D**).

Immunoblotting of individual CI subunits was carried out on isolated mitochondria from each genotype. The abundance of N-module subunits NDUFS4 (18 kD) and 75 kD increased significantly in *rmb2::ciaf1* compared to *ciaf1* (*p*<0.05, *n*=3) (**Figure 2E-F**). No significant difference in the abundance of CI membrane arm subunits such as B14.7 and CA2, CIII subunit Rieske iron-sulfur protein (RISP) or the ATP synthase subunit-β (ATPβ) (*p*<0.05, *n*=3) was identified in *rmb2::ciaf1* compared to *ciaf1* (**Figure 2E-F**). Additionally, no significant changes were observed in the abundance of the mitochondrial FTSH proteases, including the matrix-facing FTSH3 and FTSH10 and the IMS-facing FTSH4 protease in *rmb2::ciaf1* compared to *ciaf1* (**Figure 2E-F**). Translocase of the outer membrane (TOM40), an outer membrane mitochondrial protein was used as a loading control.

### *rmb2::ciaf1* exhibits restored abundances of CI matrix arm subunits

To assess global changes to the proteome in *rmb2::ciaf1* we used high resolution isoelectric focusing (HiRIEF) liquid chromatography mass spectrometry (LC-MS) proteomics of rosette leaf tissue (Branca et al., 2014). Previously published analysis of *ciaf1* indicated a decrease of CI NQ-module subunits compared to Col-0, (**Figure 3A-B, Figure S2**) (Ivanova et al., 2019). Analysis of *rmb2::ciaf1* revealed similar protein abundance profiles to what we had observed for *rmb1::ciaf1* (Ivanova et al., 2021)*. rmb2::ciaf1* and *rmb1::ciaf1* both exhibit an increased abundance of ~ 1.5-fold of matrix arm subunits (**Figure 3B-C, Figure S2**).

**Figure 3.**
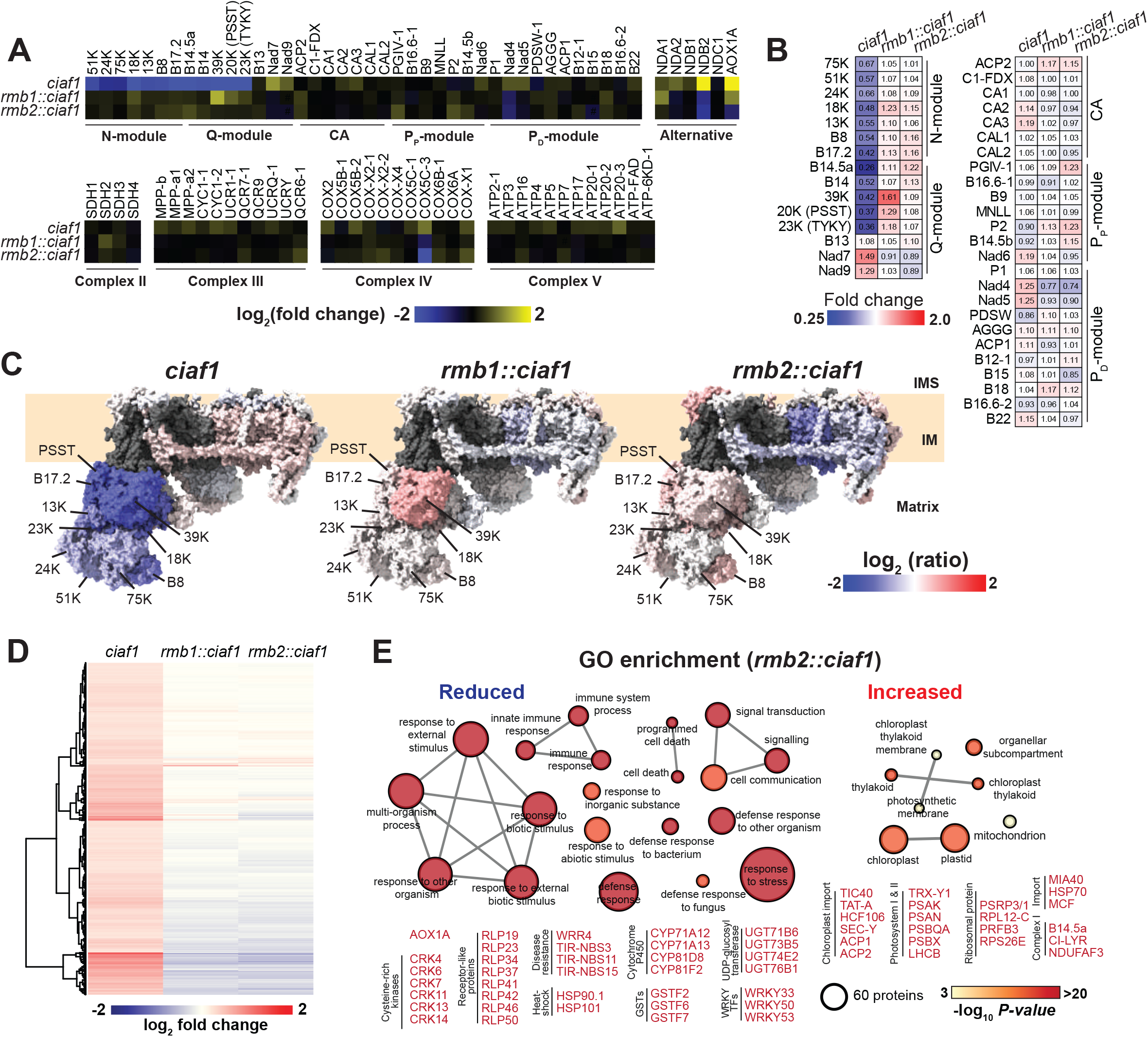
High-resolution isoelectric focusing (HiRIEF) proteomics analysis of *ciaf1, rmb1::ciaf1* (mutation in FTSH3^P415L^), and *rmb2::ciaf1* (mutation in PSST^S70F^). **A.** Heatmap showing log_2_ fold-change of protein abundance for OXPHOS subunits as measured by HiRIEF proteomics, *ciaf1* compared to Col-0, *rmb1::ciaf1* and *rmb2::ciaf1* compared to their *ciaf1* background. **B.** Relative abundance (fold-change) complex I subunits in *ciaf1, rmb1::ciaf1*, and *rmb2::ciaf1*. Complex I modules are indicated: NADH-binding (N); ubiquinone-binding (Q); carbonic anhydrase (CA); proximal proton-pumping (PP); and distal proton-pumping (PD) modules. **C.** Topographical heatmap of log_2_ relative protein abundance fitted to cauliflower complex I structure (PDB: 7A23) (Soufari et al., 2020). The dark gray regions represent subunits that were not reliably detected. Several matrix arm domain (NQ-module) subunits are indicated. IMS: intermembrane space; IM: inner membrane. **D.** Heatmap of log_2_ transformed fold-change of protein abundance showing that most up-regulated proteins in *ciaf1* are normalised or down-regulated in *rmb1::ciaf1* and *rmb2::ciaf1* compared to their *ciaf1* genetic background. **E.** Gene Ontology (GO) term enrichment analysis of down-regulated (reduced) and up-regulated (increased) proteins in *rmb2::ciaf1* compared to *ciaf1*. The fill colour represents –log_10_ *p*-value of enrichment after Bonferroni correction. Protein groups with the highest changes are indicated. Significantly reduced: cysteine-rich kinases (CRK), receptor-like proteins (RLP), white rust resistance (WRR) and toll/interleukin receptor nucleotide-binding site (TIR-NBS) disease resistance proteins, heat-shock proteins (HSP), cytochrome P450 (CYP), glutathione-S-transferase (GST), uridine diphosphate/UDP glycosyl-transferase (UGT), and WRKY transcription factors. Significantly increased: translocase of inner chloroplastic envelope (TIC), twin-arginine translocase (TAT), acyl carrier proteins (ACP), photosystem I (PSA) and photosystem II (PSB) proteins, light-harvesting chlorophyll-binding proteins (LHCB), ribosomal protein large (RPL) and small (RPS), plastid-specific ribosomal protein (PSRP), mitochondrial intermembrane-space assembly (MIA), heat-shock protein (HSP), mitochondrial carrier family (MCF) protein, and Complex I subunits B14.5a and NADH dehydrogenase::ubiquinone subcomplex assembly factor NDUFAF3, and an unidentified Complex I LYR domain-containing protein (CI-LYR).

Proteomic analysis of stress responsive proteins in *rmb2::ciaf1* and *rmb1::ciaf1* show a normalisation of abundance compared to *ciaf1* (**Figure 3D-E**) (Ivanova et al., 2021). Stress response proteins such as alternative oxidase 1A (AOX1A), cysteine-rich kinases (CRKs), receptor-like proteins (RLPs), toll/interleukin-1 receptor nucleotide-binding site (TIR-NBS) disease-resistant proteins, heat-shock proteins (HSPs), cytochrome P450s (CYPs), glutathione-S-transferases (GSTs), and WRKY transcription factors, which were up-regulated in *ciaf1* were all observed to decrease in *rmb2::ciaf1* **(Figure 3E**). The CI defect in *ciaf1* and other CI mutants generally corresponds to increased stress responsive mechanisms (Ivanova et al., 2019; Meyer et al., 2009; Dutilleul et al., 2003) and the normalisation we observe in *rmb1::ciaf1* and *rmb2::ciaf1* likely corresponds to the restoration of CI activity.

### FTSH3 and PSST are interacting proteins and mutations within the ATPase domain of FTSH3 and within the N-terminal domain of PSST inhibit this interaction

As *rmb1::ciaf1* and *rmb2::ciaf1* display similar growth phenotypes, CI abundance and activity, and proteomic profiles, we tested the ability FTSH3 and PSST to interact. To this end, we carried out co-immunoprecipitation, yeast-2-hybrid (Y2H) interaction assays, and bimolecular fluorescence complementation (BiFC) assays. Immunoprecipitations were carried out using mitochondria isolated from a transgenic line expressing the FLAG-tagged CI subunit B14.7 (B14.7^FLAG^) (Wang et al., 2012; Klusch et al., 2022). Incubation of digitonin-treated mitochondria with anti-FLAG sepharose beads resulted in B14.7^FLAG^ being precipitated and detected with both anti-B14.7 and anti-FLAG in the co-immunoprecipitation (Co-IP) sample (**Figure 4A**). Immunodetection with antibodies raised against various CI subunits, PSST, 18 kD (NDUFS4), 75 kD, NAD1, CA2, and CAL1 revealed that intact CI was precipitated accordingly (**Figure 4A**). Immunodetection with anti-FTSH3 revealed the enrichment and presence of FTSH3 within the Co-IP sample (**Figure 4A**), suggesting an interaction between FTSH3 and CI. No band could be detected using anti-FTSH10, the partner protein of FTSH3 (Piechota et al., 2010), or FTSH4 an IMS-facing AAA+ protease (Maziak et al., 2021) suggesting the interaction with CI and FTSH3 is unique (**Figure 4A**). No other OXPHOS subunits, including CII (SDH1), CIII (RISP), CIV (COX2), and CV (ATPβ), or the outer membrane TOM40 were precipitated with B14.7^FLAG^ (**Figure 4A**).

**Figure 4.**
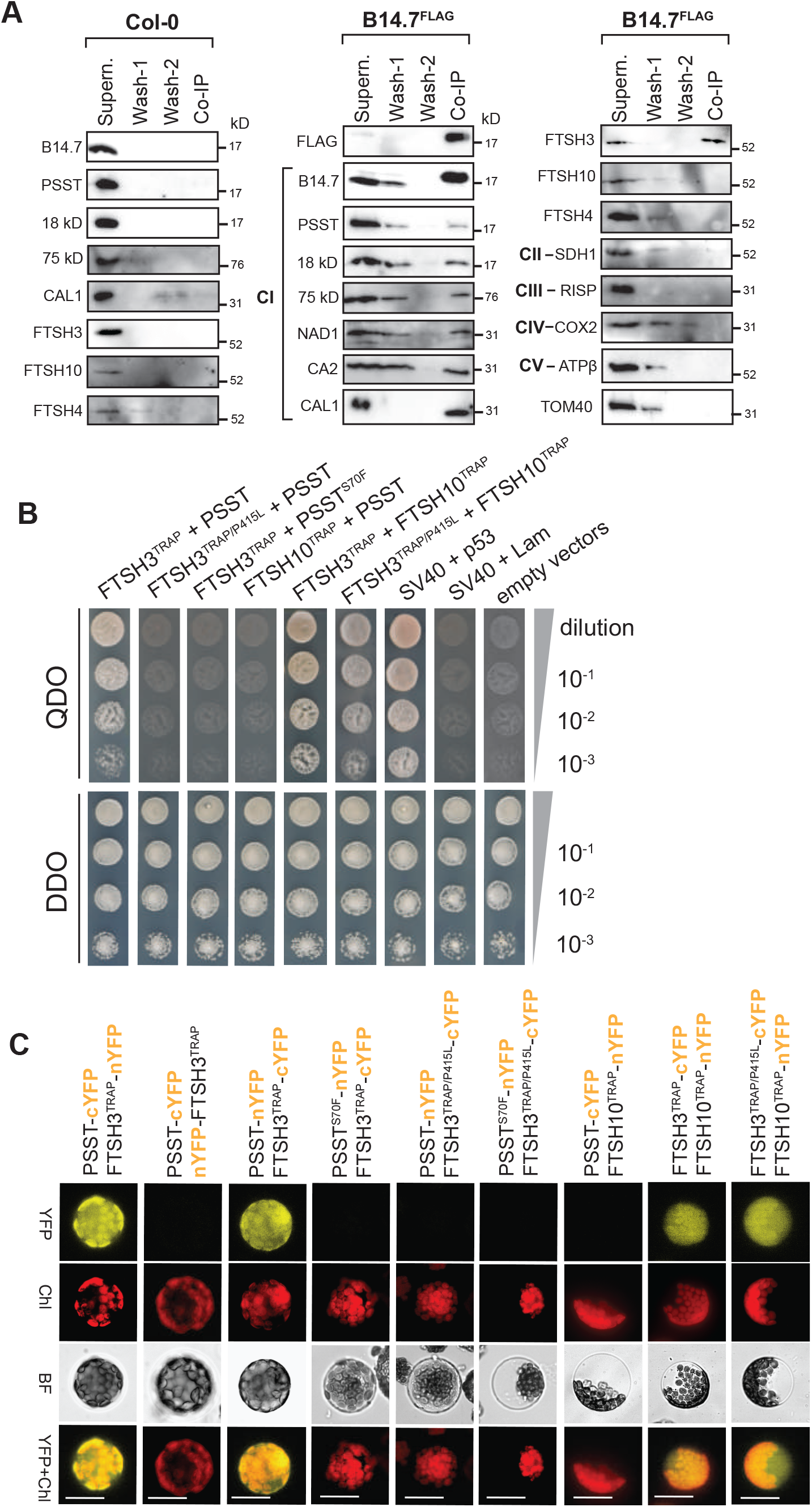
FTSH3 interacts with the complex I matrix arm subunit PSST. **A.** Co-immunoprecipitation of Columbia-0 (Col-0) and B14.7^FLAG^ mitochondria enriching complex I. The presence of immuno-detected proteins in input supernatant (supern.), first wash (wash-1), second wash (wash-2), and co-immunoprecipitated beads (Co-IP) are indicated (left), with corresponding molecular weight marker (right). **B.** Yeast-two-hybrid (Y2H) assays testing the protein interaction ability of proteolytically inactive FTSH3^TRAP^, FTSH10^TRAP^, and complex I subunit PSST, with and without FTSH3^P415L^ and PSST^S70F^ mutations. Successful diploid mating was identified by growth on double-dropout (DDO) synthetic defined (SD) media. Positive interactions were defined by growth on quadruple dropout (QDO) SD media. Serial dilutions of diploid yeast are indicated (right). Interactions between SV40 antigen with p53 and Lam were used as positive and negative controls, respectively. No interactions were observed using empty vectors. **C.** Bimolecular fluorescence complementation (BiFC) assays of proteolytically inactive FTSH3^TRAP^, FTSH10^TRAP^, and PSST, with and without FTSH3^P415L^ and PSST^S70F^ mutations. Arabidopsis mesophyll protoplasts transfected with constructs encoding proteolytically inactive FTSH3^TRAP^/ FTSH10^TRAP^ and PSST with either C-terminal or N-terminal half of yellow fluorescent protein (cYFP/nYFP). FTSH3^TRAP^ with wild-type PSST or FTSH10^TRAP^ resulted in reconstituted YFP fluorescence. Combinations involving FTSH3^P415L^ and PSST^S70F^ resulted no YFP signals, as well as between FTSH10^TRAP^ and PSST. YFP fluorescence, chlorophyll (Chl) autofluorescence, bright field (BF), and overlay of YFP and chlorophyll (YFP+Chl) are indicated. Scale bar = 20 μm.

To investigate if FTSH3 could directly interact with PSST, Y2H interaction assays were carried out using a proteolytically-inactive FTSH3 (FTSH3^TRAP^). Yeast strains transformed with FTSH3^TRAP^ and PSST were mated and tested for their ability to grow on quadruple drop out media (QDO) and double drop out media (DDO). FTSH3^TRAP^ and PSST diploid yeast grew on QDO indicating a protein-protein interaction (**Figure 4B**). A similar mating, but with the *rmb1* mutation (P415L) within FTSH3 (FTSH3^TRAP/P415L^) did not result in any growth on QDO, similarly when the *rmb2* mutation was introduced within PSST (PSST^S70F^), no interaction was observed with FTSH3^TRAP^ **(Figure 4B**). As FTSH3 has the ability to form hetero-complexes with FTSH10, the ability of FTSH10^TRAP^ to interact was PSST was also tested. No growth could be observed on QDO (**Figure 4B**). To test the interactive ability of FTSH3 and FTSH10, FTSH3^TRAP^ and FTSH10^TRAP^ were mated and grown on QDO resulting in positive growth as expected. To test if the P415L mutation within FTSH3 altered this interaction ability, FTSH3^TRAP/P415L^ and FTSH10^TRAP^ were mated and growth was observed on QDO suggesting that the P415L mutation within FTSH3 does not alter the ability of these proteins to interact (**Figure 4B**). All diploid yeast were observed to grow on DDO media confirming successful mating, alongside the positive (SV40+p53) and negative (SV40+Lam) controls **(Figure 4B**).

To confirm these protein-protein interactions *in planta*, bimolecular fluorescence complementation (BiFC) was carried out using Arabidopsis protoplasts. Co-transformation of PSST-cYFP and FTSH3^TRAP^-nYFP exhibited reconstituted fluorescence (**Figure 4C**). Whilst co-transformation of PSST-cYFP and nYFP-FTSH3^TRAP^ did not display reconstituted fluorescence (**Figure 4C**). Co-transformation of PSST-nYFP and FTSH3^TRAP^-cYFP similarly exhibited reconstituted cytosolic fluorescence indicating protein interactions, whilst co-transformation with either or both the mutated PSST (PSST^S70F^) and FTSH3^TRAP/P415L^ did not **(Figure 4C**). Co-transformation of PSST-cYFP with FTSH10^TRAP^-nYFP did not result in any reconstituted fluorescence supporting the results obtained in the Y2H assays. Additionally, FTSH3^TRAP^ and FTSH3^TRAP/P415L^ resulted in reconstituted fluorescence when co-transformed with FTSH10^TRAP^ (**Figure 4C**).

### Mutations within the ATPase domain of FTSH3 and within the N-terminal domain of PSST slows the rate of protein turnover for some CI subunits

Disruption of mitochondrial AAA+ proteases has resulted in altered turnover rates for various OXPHOS subunits (Li et al., 2017; Szczepanowska et al., 2020). To assess protein turnover, we carried out stable isotope ^15^N-labelling of Col-0, *ciaf1, rmb1::ciaf1* and *rmb2::ciaf1* seedlings, followed by mitochondrial isolation and quadrupole time-of-flight mass spectrometry (QTOF-MS) of soluble matrix fractions resulting in the detection of protein turnover rates for 832 out of 1547 mitochondrial proteins (**Table S3**). Degradation or turnover rate (K_D_) of individual proteins was measured based on fold-change of protein abundance (FCP) of ^15^N-labelled and unlabelled proteins over a period of labelling time using an equation stipulated previously (Li et al., 2017). Analysis of degradation rates (K_D_) of CI subunits from *ciaf1* compared to Col-0, show overall faster turnover rates for subunits located within the CI matrix NQ-module (**Figure 5A**). This is consistent with the previous finding that the NQ-module subunits are less abundant due to the incorrect assembly and maturation in *ciaf1* (Ivanova et al., 2019). No notable changes in turnover rates could be identified for other OXPHOS subunits (Ivanova et al., 2019) (**Figure S3**).

**Figure 5.**
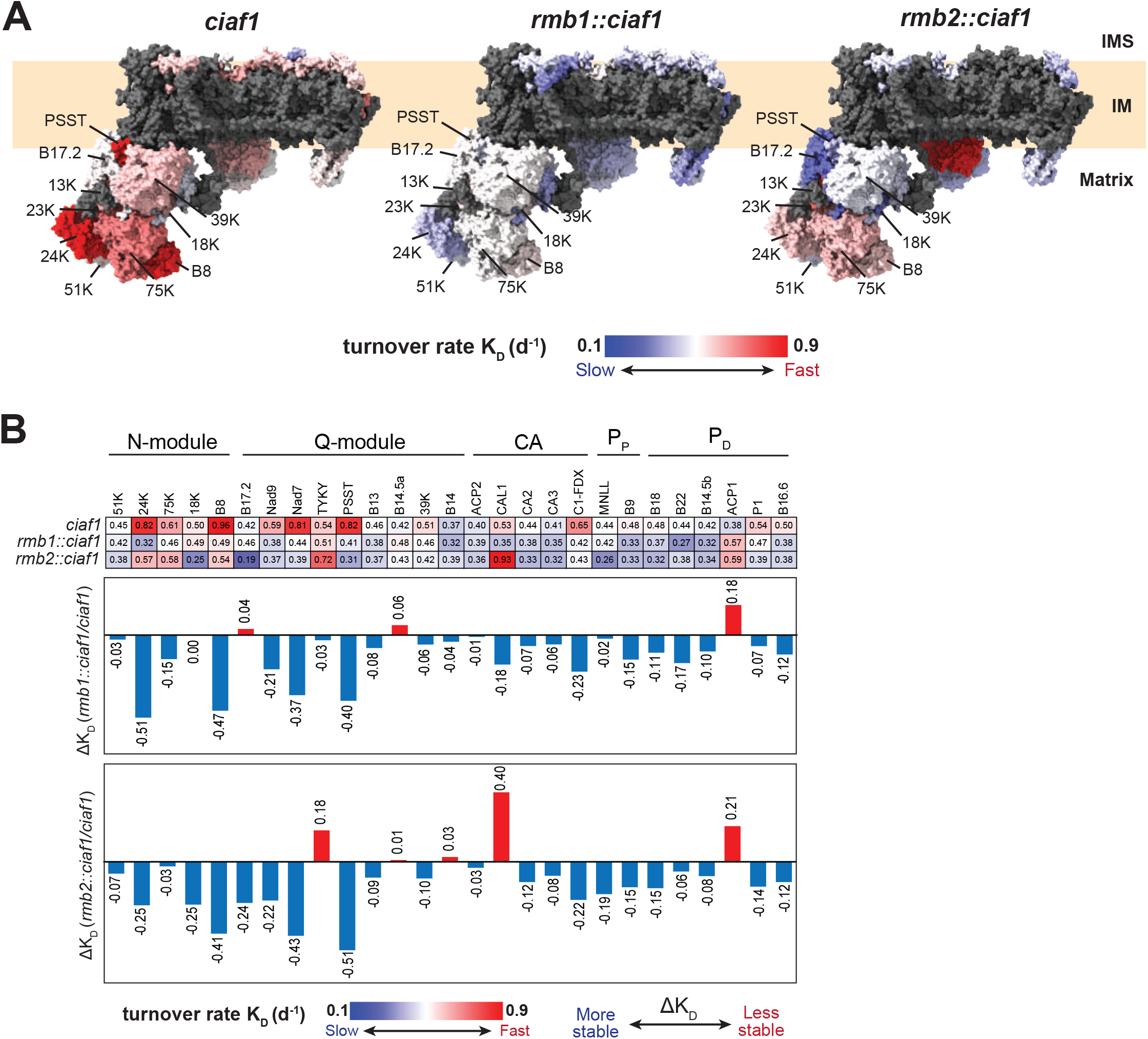
Mutations in *rmb1::ciaf1* (FTSH3^P415L^) and *rmb2::ciaf1* (PSST^S70F^) displayed slowed turnover rates of complex I subunits. **A.** Topographical heatmap of protein turnover rate K_D_ (d^-1^) measured from matrix-soluble fraction fitted into complex I structure (PDB: 7A23) (Soufari et al., 2020). Heatmap shows subunits with relatively slow (blue, KD = 0.1) to relative fast turnover rate (red, K_D_ ≥ 0.9) (Li et al., 2017). Several subunits are indicated. Dark gray region represents subunits that were unreliably detected. **B.** Comparison of turnover rates K_D_ (d^-1^) for individual complex I subunits in *ciaf1, rmb1::ciaf1*, and *rmb2::ciaf1* shows subunits with relatively slow (blue, K_D_ = 0.1) to relative fast turnover rate (red, K_D_ ≥ 0.9). Change in turnover rate (ΔK_D_) showing the difference of turnover rate of *rmb1::ciaf1* and *rmb2::ciaf1* compared to *ciaf1* background, with negative value (blue) indicates more stable subunits and positive value (red) represents less stable subunits. Specific complex I modules are indicated: NADH-binding (N); ubiquinone-binding (Q); carbonic anhydrase (CA); proximal proton-pumping (PP); and distal proton-pumping (PD) modules.

Analysis of turnover rates from *rmb1::ciaf1* compared to *ciaf1* indicate slower degradation rates (K_D_) of NQ-module subunits confirming a slowing of disassembly and turnover (**Figure 5A)**. Comparison of turnover rates (ΔK_D_), which is the difference of turnover rate between EMS mutants *rmb1::ciaf1* and *rmb2::ciaf1* and background genotype *ciaf1*, showed that on average, CI subunits in both *rmb1::ciaf1* and *rmb2::ciaf1* have ΔK_D_ of −0.12 d^-1^ compared to *ciaf1* (**Figure 5B**), which corresponds to an increased half-life of 6 days on average (**Figure 5B**). The mutation within the N-terminal domain of PSST (S70F) results in relatively slower turnover rates (ΔKD) of CI subunits and consequently a more stable CI compared to *ciaf1* **(Figure 5A).** The effect is more notable in the NQ module subunits, such as PSST, B8, Nad7, Nad9, and 24 kD, which exhibit slower turnover rates by ~1.5-2 fold compared to *ciaf1* (**Figure 5B**). In *ciaf1*, PSST has a relatively fast turnover rate (0.82 d^-1^, half-life 20.4 h) whilst in *rmb1::ciaf1* and *rmb2::ciaf1*, PSST has substantially slower turnover rates of 0.41 d^-1^ (half-life 40 h) and 0.31 d^-1^ (half-life 54 h) respectively (**Figure 5B**). The overall trend of slowed turnover rates in *rmb1::ciaf1* and *rmb2::ciaf1* suggests that mutations within FTSH3 or PSST independently slow the turnover of CI subunits resulting in an accumulation of CI and a partial restoration of plant growth phenotypes (**Figure 1, 3**).

### The *rmb2::ciaf1* mutation within PSST is likely to affect the N-terminal domain structure

The consequences of both *rmb1::ciaf1* and *rmb2::ciaf1* mutations were examined at the molecular level. As plant FTSH3 structures are not available, we modelled Arabidopsis FTSH3 using the human AFG3L2 structure as a template (PDB ID: 6NYY) (Puchades et al., 2019) (**Figure S4**). AFG3L2 is a matrix-facing AAA+ (*m*-AAA) protease closely related to bacterial FTSH and plant *m*-AAA proteases. AFG3L2 exhibits 64.3% similarity and 49.0% identity to Arabidopsis FTSH3, and contains the conserved ATPase and protease domains (**Figure S4**). The ATPase domain of each AFG3L2 monomer contains a ATP-binding pocket that powers the movement of two substrate-recognising loops called pore-loop 1 and pore-loop 2 to translocate substrates into the proteolytic domain (Puchades et al., 2017, 2019) (**Figure 6A**). AFG3L2 consists of homo-hexameric ATPase and protease domains, and a peptide substrate moiety (Puchades et al., 2019) allowing PSST to be modelled as a putative substrate for FTSH3 homo-hexamer, based on the previously determined plant CI structure (PDB ID: 7A23) (Soufari et al., 2020) (**Figure S5**). The substrate-bound structure shows that PSST can be accommodated into the FTSH3 AAA+ pore and interact with the substrate-recognising pore-loop 1 (**Figure 6A**). Modelling suggests the N-terminal peptide of PSST has a diameter of 9.01 Á and displays no unfavourable clash with substrate-recognising phenylalanine residues (F395) (Puchades et al., 2019) (**Figure 6B**). However, modelling of PSST harbouring the S70F mutation suggests a larger helix size (11.50 Á diameter), due to the bulky benzyl group of phenylalanine having the potential for unfavourable interactions with F395 (**Figure 6B**). This may lead to a thermodynamic instability (Galemou Yoga et al., 2019), preventing an interaction between FTSH3 and PSST^S70F^.

**Figure 6.**
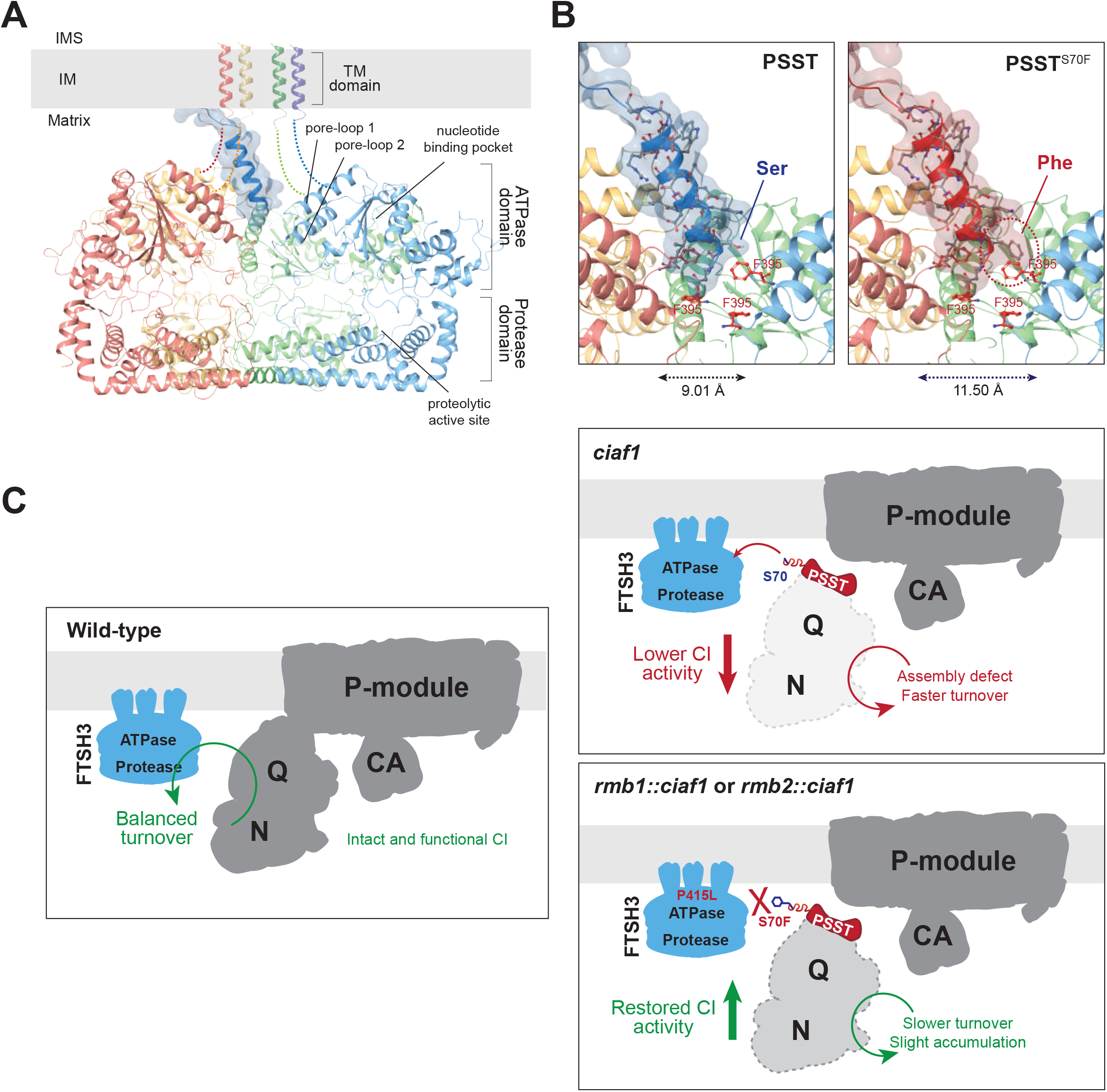
Interaction of FTSH3 and PSST allows disassembly and degradation of complex I NQ-module. **A.** Homology model structure of FTSH3 (multi-coloured ribbons) and the first 30-aa of PSST representing the N-terminal domain (dark blue ribbon and surface). The interaction was modelled based on human *m*-AAA FTSH homologue AFG3L2 with a bound peptide substrate (PDB: 6NYY) (Puchades et al., 2019). **B.** FTSH3-PSST molecular interaction analysis based on homology-modelled structure. PSST with 9.01 Å helical diameter (blue ribbon) fitted to FTSH3 AAA+ pore in the ATPase domain. Substrate-recognising phenylalanine residues (F395) situated in pore-loop 1 from different monomers are indicated (red sticks). EMS-mutated *rmb2::ciaf1* PSST (PSST^S70F^) has a wider overall diameter (11.50 Å) (red ribbon), resulted in an unfavourable clash with one of F395 residues (red circle). **C.** Proposed model of FTSH3-mediated regulation of complex I matrix arm domain in wild-type, *ciaf1, rmb1::ciaf1* and *rmb2::ciaf1*. PSST binding by FTSH3 allows its proteolytic activity against complex I matrix arm NQ-module. Assembly defect of NQ-module in *ciaf1* induces its unfolding, disassembly, and subsequent degradation, leading to lower abundance and complex I overall activity. Mutation of either FTSH3 ATPase domain in *rmb1::ciaf1* (FTSH3^P415L^) and N-terminal domain of PSST in *rmb2::ciaf1* (PSST^S70F^) abolished their interaction, resulted in slower turnover and slight accumulation of NQ-module subunits, which partially restored complex I activity.

## Discussion

Here, we uncover the mechanism for the regulation of CI turnover by the inner membrane bound FTSH3 protease, via an interaction with the CI matrix arm subunit PSST. Using a CI defective mutant *ciaf1*, we generated two independent EMS-induced mutants that displayed restored CI activity and plant growth. We demonstrated that the two independent mutations, one within the ATPase domain of FTSH3^P415L^ (*rmb1::ciaf1*) (Ivanova et al., 2021) and another within the N-terminal domain of PSST^S70F^ (*rmb2::ciaf1*) were responsible for restoration of CI in *ciaf1*. Furthermore, protein interaction assays show that FTSH3, and not FTSH10 interacts with PSST. Mutations within the FTSH3 ATPase domain (FTSH3^P415L^) or the N-terminal domain of PSST (PSST^S70F^) abolished this interaction, demonstrating the importance of these residues and domains for substrate recognition and subsequent NQ-module disassembly and proteolytic degradation. These mutations resulted in slowed protein turnover rates of NQ-module subunits were slowed compared to *ciaf1* resulting in an increased accumulation of the NQ-module subunits, and ultimately increased CI abundance and activity (**Figure 6C**).

We previously showed that complementation of *rmb1::ciaf1* with FTSH3 resulted in a reverted phenotype typical of *ciaf1*, i.e. small and developmentally delayed. Complementation using a proteolytic inactive FTSH3 similarly reverted the phenotype indicating that the proteolytic function of FTSH3 is not required for the restoration of CI abundance in this case (Ivanova et al., 2021). Furthermore, deletion of FTSH3 in a Col-0 background has no observable effect to CI (Kolodziejczak et al., 2018; Ivanova et al., 2021) under normal conditions. In this case, FTSH3 activity is unlikely to be compensated by FTSH10. Although FTSH3 can form a hetero-hexamer with FTSH10 (Piechota et al., 2010). FTSH10 was not detected in our CI immunoprecipitations (**Figure 4A**). Further, no interaction of FTSH10 with PSST was observed using Y2H and BiFC assays (**Figure 4C-D**). It is likely that a FTSH3 homo-hexamer and not a FTSH3/FTSH10 hetero-hexamer is involved in PSST recognition and disassembly of CI matrix arm. A similar situation has been previously reported with the human FTSH3 ortholog AFG3L2. AFG3L2 assembles as a homo-hexamer, or as hetero-hexamer with paraplegin (SPG7), a AAA+ protease (Puchades et al., 2019; Koppen et al., 2007). However, only the AFG3L2 homo-hexamer is responsible for maturation and degradation of OXPHOS components (Arlt et al., 1998; Koppen et al., 2007).

Previous studies in mammals have shown that damaged CI subunits are disassociated, degraded, and replaced with newly synthesised subunits to maintain CI activity (Szczepanowska et al., 2020). In plants, the NQ-module subunits display relatively higher turnover rates compared to the membrane arm domain subunits and other mitochondrial OXPHOS subunits (Nelson et al., 2013; Li et al., 2017). We previously showed that under a CI assembly defect, FTSH3 plays a role in the disassembly of the NQ module domain likely via its ATPase domain (Ivanova et al., 2021), required for substrate recognition and ATP-dependent protein unfolding, as previously observed in other AAA+ proteases (Puchades et al., 2017, 2019; Shin et al., 2020, 2021). In this study we uncover the mechanism of how this occurs. We identified PSST as a substrate of FTSH3. PSST, an orthologue of human NDUFS7, is a 20 kD Q module subunit that is located within the interface of the matrix and membrane arms domain and is conserved across species (Fiedorczuk et al., 2016; Parey et al., 2019; Soufari et al., 2020; Klusch et al., 2021; Baradaran et al., 2013; Kolata and Efremov, 2021) (**Figure S4**). Mutation of PSST conserved loop residues in aerobic yeast *Yarrowia lypolitica* resulted in reduced electron transfer to ubiquinone (Galemou Yoga et al., 2019), whilst deletion of PSST in Arabidopsis results in severe developmental delays confirming its essential function (Kühn et al., 2015). PSST is located adjacent to the membrane-bound P_P_ module, forming the ubiquinone-binding tunnel at the interface of the Q module and Nad7 subunit in the P_P_ module (Soufari et al., 2020; Klusch et al., 2021). The N-terminal domain of PSST is exposed to the matrix, particularly when the NQ module is not correctly assembled to the holocomplex (**Supplementary Figure 5**), suggesting that PSST is accessible for recognition by FTSH3 for unfolding and subsequent proteolysis (**Figure 6C**).

Substrate unfolding by AAA+ proteases has previously been determined in other species. Human YME1L, a FTSH4 ortholog recognises its substrate via an accessible N-terminal sequence (Shi et al., 2016). In addition, the accessibility of the substrate peptide through the AAA+ pore is required for protease activity (Shi et al., 2016). The protein unfolding activity of AAA+ proteases depends on ATP hydrolysis and the size of the unfolded polypeptide substrate (Martin et al., 2008). Homology modelling of the S70F mutation in PSST indicates an increased diameter of the helix (**Figure 6B**), potentially resulting in inaccessibility through the FTSH3 AAA pore and altering the binding energy of a FTSH3-PSST interaction (**Figure 4**).

A similar FTSH protease-mediated turnover mechanism has been proposed for photosystem II (PSII) repair cycle, where the reaction centre D1 protein is directly regulated by FTSH protease homologues. The light-absorbing activity of D1 results in its subsequent photo-damage and rapid turnover (Silva et al., 2003). Loss of the thylakoid membrane-associated FTSH2 and FTSH5 resulted in an accumulation of damaged D1 proteins (Kato et al., 2009). In the cyanobacterium *Synechocystis* sp PCC 6803, a FTSH homologue (slr0228) was found to have a direct role in the binding, unfolding, and degradation of D1 during the early stage of PSII repair cycle (Silva et al., 2003).

In mammals, CLPXP, another AAA+ protease was shown to actively degrade matrix-facing N-module domain subunits (Szczepanowska et al., 2020). Immunoprecipitations identified that the matrix located CLPXP interacted specificity with the matrix-facing NQ subunits NDUFS1, NDUFV1, and NDUFV2 (orthologous to plant 75 kD, 51 kD, and 24 kD, respectively) (Szczepanowska et al., 2020; Subrahmanian et al., 2016). Similarly, in Arabidopsis, knockdown of the CLPXP subunit, CLPP2 resulted in an accumulation of the 75 kD, 51 kD, and 24 kD subunits and an accumulation of a matrix submodule containing the 24 kD and 51 kD subunits, (Petereit et al., 2020) suggesting a role in the proteolysis of CI.

In comparison to other AAA+ proteases, FTSH homologues exhibit weak protein unfolding activity when measured by their efficiency in converting ATP hydrolysis to protein degradation (Shi et al., 2016). Indeed in bacteria, FTSH was also proposed to exhibit weak unfoldase activity (Herman et al., 2003; Kihara et al., 1999). Rather, bacterial FTSH recognised substrates when misfolded or disassembled from its complex, allowing it to be unfolded and degraded. Therefore, FTSH regulates the abundance of membrane-bound proteins depending on their folding state or their association to a complex (Kihara et al., 1995, 1999; Herman et al., 2003) allowing FTSH proteases to recognise damaged proteins in membrane-bound complexes (Herman et al., 2003). Our identification of a disassembly function of FTSH3 was only identified in a dysfunctional CI mutant (Ivanova et al., 2020), which is consistent with the proposal that FTSH proteases recognise unfolded substrates.

In conclusion, here we have identified and demonstrated the mechanism of CI turnover by FTSH3. Through its interaction with PSST, FTSH3 plays a role in the disassociation of the NQ module from holo-CI, allowing for selective degradation and turnover of damaged and non-functional subunits.

## Materials and Methods

### Plant material, growth, and phenotyping

All plants used in this study were of Columbia-0 (Col-0) ecotype background. Seeds were surface-sterilised with chlorine gas and sown on half-strength Murashige-Skoog (½ MS) liquid or agar medium (pH 5.7) supplemented with 1.5% (w/v) sucrose and 0.1% (v/v) Gamborg B5 vitamins. A peat moss, vermiculite, and perlite (3:1:1) mix was used for soil growth. All plants were grown under long-day photoperiod (16h light, 8h dark) with 100 μmol/m^2^s light intensity at 22°C.

### Forward genetics and whole genome sequencing

EMS treatment was used to mutagenize the CI assembly factor 1 knockdown mutant *ciaf1* (SALK_143656) as previously described (Weigel and Glazebrook, 2006; Ivanova et al., 2021). The progeny of approximately 2,000 individual M1 plants were grown on ½ MS agar media and visually screened to identify plants exhibiting substantial increased growth compared to *ciaf1* (Ivanova et al., 2019). Candidate lines were confirmed in the M3 generation following three rounds of back-crossing to *ciaf1*. Six homozygous *rmb2::ciaf1* lines from the segregating back-crossed population were used for sequencing analysis.

Genomic DNA was isolated using DNeasy Total DNA Purification Maxi Kit (Qiagen) according to manufacturer’s instructions. Sequencing libraries were generated using Nextera DNA Flex Library Prep Kit (Illumina) from total genomic DNA and sequenced with a 70-bp read length on a NextSeq 500 system (Illumina). Whole genome sequencing was done using the HiSeq 2500 (Illumina). Sequencing reads were aligned to the Arabidopsis reference genome (Araport11) using bowtie2 v2.3.4.1 (Langmead and Salzberg, 2012). Mutations were identified using the “mpileup” and “call” commands of bfctools v1.4 with default parameters (http://www.htslib.org/doc/bcftools.html). A single nucleotide mutation C to T in chromosome 5 within At5g11770 was present in all six homozygous revertant lines.

### Cloning, plasmid constructs, and line generation

The PSST (At5g11770), B14.7 (At2g42210) and FTSH10 (AT1G07510) protein-encoding sequence (CDS) were cloned from Col-0 complementary DNA (cDNA). Cloning was carried using Gateway^®^ site-specific recombination (Invitrogen) (Katzen, 2007) into the pDONR207 entry vector using gene-specific primers (**Table S1**). Site-directed mutagenesis was performed using QuikChange™ kit (Stratagene) to generate FTSH3^P415L^, PSST^S70F^, FTSH3^TRAP^ (H586G) and FTSH10^TRAP^ (H586G) with primers listed in **Table S1.** Clones were recombined into: pH2GW7 (with CaMV 35S promoter) for over-expression lines (Karimi et al., 2002), pGWB11 for C-terminal FLAG tagged B14.7 (Nakagawa et al., 2007), pGBKCg/pGBKT7 and pGADCg/pGADT7g for yeast-2-hybrid assays (Stellberger et al., 2010), pSAT4-DEST-nEYFP-C1 (pE3136), pSAT4-DEST-nEYFP-N1 (pE3134), pSAT5-DEST-cEYFP-C1 (pE3130), and pSAT5-DEST-cEYFP-N1 (pE3132) for BiFC assays (Citovsky et al., 2006). The full-length PSST gene including the 1 kb upstream promoter region was cloned into pCAMBIA1300 using NEBuilder^®^ HiFi DNA assembly master mix (NEBioLabs) from genomic DNA. Complementation and over-expression lines were generated using *Agrobacterium tumefaciens* via the floral dip method (Karimi et al., 2002).

### Reverse-Transcription quantitative PCR (RT-qPCR)

Total RNA from 14d old agar-grown seedlings was isolated using FavorPrep total RNA purification mini kit (Favorgen). cDNA was generated using a high-capacity cDNA reverse transcription kit (Applied Biosystem). RT-qPCR standards were generated from Col-0 cDNA pool using MyTaq polymerase mix (Bioline) and purified using Favorprep PCR Purification Kit (Favorgen) according to manufacturer’s instructions and listed primers (**Table S1**). RT-qPCR was carried out in LightCycler 480 (Roche) using SYBR Green master mix (Roche) and gene-specific primers (**Table S1**). *ACTIN2* (At3g18780) was used as a control. All measurements were done from at least three independent biological and two technical replicates.

### Bimolecular fluorescence complementation (BiFC) assay

BiFC assays were carried out using transiently transformed isolated Arabidopsis leaf mesophyll protoplasts as previously described (Yoo et al., 2007; Wu et al., 2009). Briefly, 12-15 fully expanded 3–4-week-old rosette leaves were harvested and sandwiched with tape to remove the lower epidermis layer. Stripped leaves were incubated in freshly prepared enzyme solution (0.5% w/v cellulase, 0.25% w/v pectolyase, 0.4 M mannitol, 20 mM KCl, 10 mM CaCl_2_, 20 mM methyl ethane sulfonate (MES) pH 5.7, 0.1% w/v bovine serum albumin (BSA), sterilised through a 0.45 μm syringe filter) for 1h at 40 rpm in the light. Protoplasts were harvested by centrifugation at 100*g* 4°C for 12 min, washed twice with W5 solution (154 mM NaCl, 125 mM CaCl_2_, 5 mM KCl, 5 mM glucose, 2 mM MES pH 5.7), and incubated on ice for 30 min in the dark. Protoplast concentration was adjusted to ~10^5^ cells/mL in 2 mL MMG solution (0.4 M mannitol, 15 mM MgCl_2_, 5 mM glucose, 4 mM MES pH 5.7, sterilised through a 0.45 μm syringe filter). To transform protoplasts, 10 μg of plasmid (in a 10 μL total volume) and 110 μL freshly prepared sterile polyethylene glycol (PEG) solution (40% w/v PEG, 0.1 M CaCl_2_, 0.2 M mannitol) were added to 500 μL protoplast. The mixture was incubated in the dark for 20 min, washed twice with 440 μL W5 solution (centrifuged 100 *g* for 2 min), and resuspended in 500 μL W5 solution. The transformed mixture was incubated overnight (12h) in the dark before visualised in BX61 fluorescence microscope (Olympus). Fluorescence was visualised with the excitation/emission wavelengths of 540-550/565-575 nm for chloroplast autofluorescence (TRITC) and 510-520/525-535 nm for enhanced yellow fluorescent protein (eYFP).

### Yeast-2-hybrid (Y2H) assay

Y2H assays were carried out as previously described (Ivanova et al., 2019). Bait constructs containing DNA-binding domain/DBD (pGBKCg/PGBKT7) (Stellberger et al., 2010) were transformed into the Y187 yeast strain, while prey constructs containing the activating domain/AD (pGADCg/pGADT7) (Stellberger et al., 2010) were transformed into the AH109 strain. Mating was carried out in a 96-well plate according to manufacturer’s instructions (Matchmaker GAL4 Two Hybrid System (Clontech/TakaraBio). Mated diploid strains were serially diluted and plated on - Leu–Trp double dropout (DDO) and –Leu–Trp–His–Ade quadruple dropout (QDO) synthetic defined (SD) agar media and incubated for 4d at 30°C. Interactions were considered positive if growth was observed on QDO in all three biological replicates.

### Mitochondrial isolation, gel electrophoresis, and immunoblotting

Mitochondria were isolated from 14d-old seedlings grown in liquid ½ MS media pots with 200 rpm shaking as described previously (Duncan et al., 2017a). For blue native polyacrylamide gel electrophoresis (BN-PAGE), samples was prepared by solubilising mitochondria with 5% (w/v) digitonin as previously described (Eubel et al., 2005). Samples were resolved using precast NativePAGE™ 4-16% Bis-Tris gels (Invitrogen). CI in-gel activity staining was carried out as previously described (Schertl and Braun, 2015). Protein separation and identification was done by sodium dodecyl sulfate polyacrylamide gel electrophoresis (SDS-PAGE) (4-12% Bis-polyacrylamide gel) of solubilised intact mitochondria followed by immunoblotting. After SDS-PAGE, proteins were transferred onto PVDF membrane and immunodetection carried out using antibodies: 75 kD (PhytoAB), NDUFS4 (18 kD) (Meyer et al., 2009), PSST (20 kD) (Meyer et al., 2011), B14.7 (Wang et al., 2012), NAD1 (PhytoAB), CAL1 (Fromm et al., 2016), CA2 (Perales et al., 2005), SDH1 (Peters et al., 2012), RISP (Carrie et al., 2010), COXII (Agrisera), ATPβ (Agrisera), TOM40 (Carrie et al., 2010), TIM17-2 (Murcha et al., 2003), FTSH3 (Kolodziejczak et al., 2007), FTSH4 (Urantowka et al., 2005), and FTSH10 (Piechota et al., 2010). For densitometric measurement of band intensity, blots were analysed in ImageJ (Schneider et al., 2012).

### Immunoprecipitation

Mitochondria were isolated from B14.7^FLAG^ tagged lines. Anti-FLAG (anti-DYKDDDK) monoclonal M2 sepharose beads (Sigma Aldrich) was used to bind and precipitate B14.7^FLAG^ along with any associated proteins. Freshly isolated mitochondria were ruptured with 5% (w/v) digitonin in 0.5 mL lysis buffer containing 20 mM Tris-Cl, 50 mM NaCl, 0.1 mM EDTA, 10% (v/v) glycerol, and 1 mM phenylmethanesulfonyl fluoride (PMSF), and 10 μL of protease inhibitor (protease and phosphatase inhibitor cocktail dissolved in 1 mL water) (Sigma Aldrich), and incubated on ice for 30 min.

The lysed mitochondrial mixture was centrifuged at 20,000 *g* for 15 min at 4°C. The supernatant was removed and added to 150 μL of prewashed anti-FLAG tag beads and incubated for 16 h at 4°C on a rotary shaker. The supernatant was removed by centrifugation at 1,000 *g* for 1 min followed by washing the beads in 1 mL 1X PBS buffer (140 mM NaCl, 10.1 mM Na_2_HPO4, 1.76 mM KH_2_PO_4_ and 2.68 mM KCl) three times. Supernatants, wash fractions and beads were stored at −80°C prior to SDS-PAGE and immunoblotting.

### Complex I activity assay

CI activity (NADH:ubiquinone reductase) was measured as previously outlined (Huang et al., 2015). The decrease of *A*_420_ was recorded after adding 1 μg mitochondrial sample to 200 μL of reaction solution containing 50 mM Tris-HCl buffer pH 7.2, 50 mM NaCl, 0.2 mM deamino-NADH, and 1 mM [Fe(CN)_6_]^3-^ (ε 1.03 mM^-1^ cm^-1^ at 420 nm). CI activity was determined as the rate of [Fe(CN)_6_]^3-^ reduction as it received electrons from NADH oxidation by CI.

### HiRIEF proteomics analysis

Steady-state proteomics of Col-0, *ciaf1-1, rmb1::ciaf1* and *rmb2::ciaf1* lines were carried out using high-resolution isoelectric focusing (HiRIEF) followed by liquid chromatography mass spectrometry (LC-MS) (Branca et al., 2014) as described previously (Ivanova et al., 2019). The heatmap of significant changes was generated in R using ggplot2 package with Euclidian distribution and Ward’s hierarchical clustering method. Gene Ontology (GO) term enrichment was carried out using ClueGO plugin in Cytoscape (Bindea et al., 2009; Shannon et al., 2003), with two-sided hypergeometric enrichment and Bonferroni *p*-value correction. Relative abundance of proteins were mapped to CI structure from cauliflower (*Brassica oleracea var. bortyris;* PDB: 7A23) (Soufari et al., 2020). Protein structures were visualised in UCSF ChimeraX (Pettersen et al., 2021), using datasets available in **Table S2**.

### Proteomics analysis for turnover rate

Heavy nitrogen labelling of Arabidopsis seedlings was carried out as previously described (Li et al., 2017). Briefly, seeds were grown in 80 mL of liquid ½ MS medium for 10 d. The seedlings were washed with sterile liquid ½ MS media without nitrogen for three times before replaced with modified ^15^N growth medium containing ½ MS medium without nitrogen, 0.96 g/L ^15^NH_4_^15^NO_3_, and 0.83 g/L K^15^NO_3_ and grown for another 4 d. The 14 d old seedlings were harvested and mitochondria isolated as described previously (Duncan et al., 2017b). Matrix fractions were collected from mitochondrial samples (corresponding to 500 μg protein content) by 3× freeze-thaw cycle (20 min at −20°C, followed by 20 min at 4°C), vortexed at 4°C, and then centrifuged at 20,000 *g* for 30 min at 4°C. Trypsin digestion, identification, and quantification of peptides using mass spectrometry were done as previously described (Li et al., 2017).

### Sequence alignment and structural homology modelling

FTSH3, FTSH10, and PSST homologue sequences were retrieved from Uniprot (https://www.uniprot.org/) (Bateman et al., 2017). Sequence alignments were carried out using Clustal Omega (https://www.ebi.ac.uk/Tools/msa/clustalo/) (Sievers et al., 2011). Arabidopsis FTSH3 structure was modelled from full protein sequence retrieved from TAIR (https://www.arabidopsis.org/) (Huala et al., 2001). Homology modelling was carried out using Phyre2 (Kelley et al., 2015), with human m-AAA protease AFG3L2 structure (PDB: 6NYY) as the reference template (Puchades et al., 2019). PSST N-terminal domain structure was retrieved from the cauliflower CI structure (PDB: 7A23) (Soufari et al., 2020). Structural visualisation, surface generation, structural superimposition, and *in silico* point mutations were carried out in Maestro (Schrödinger, LLC) and ChimeraX (Pettersen et al., 2021).

### Statistical analysis

All replicates stated in this study are biological replicates, unless stated otherwise. Statistical difference between multiple means was determined using single-factor ANOVA (95% confidence level, *p*<0.05). Significantly different means were then determined using Tukey-Kramer post-hoc honest significant test.

### Data availability

Proteomics data used in this study have been deposited publicly online in the ProteomeXchange Consortium via the Proteomics Identification Database (PRIDE) partner repository (https://www.ebi.ac.uk/pride/) with a project identifier PXD011795. Proteomics data of *ciaf1* and *rmb1::ciaf1* has been published in our previous studies (Ivanova et al., 2019, 2021). Protein abundance data of Col-0 (WT), *ciaf1, rmb1::ciaf1*, and *rmb2::ciaf1* can be accessed in the annotated protein table as Col0, ciaf1-1, Mut1, and Mut3, respectively. Sequence data associated with this study can be found in The Arabidopsis Information Resource (TAIR) database (https://www.arabidopsis.org/) with the following Arabidopsis Genome Initiative (AGI) identifier: AT3G18780 (Actin2), AT5G08670 (ATPβ), AT2G42210 (B14.7), AT5G63510 (CAL1), AT3G48680 (CAL2), AT1G47260 (CA2), AT1G76060 (CIAF1), AT2G29080 (FTSH3), AT5G67590 (18 kD/NDUFS4), AT5G37510 (75 kD), AT5G11770 (20 kD/PSST), AT3G20000 (TOM40), AT3G01280 (VDAC1/porin), AT1G07510 (FTSH10), AT2G26140 (FTSH4), At5G13430 (RISP), AT5G66760 (SDH1-1), and ATMG00160 (COX2).

## Author contribution

A.S.G, A.I and M.W.M performed experiments and analysed the data. A.S.G prepared all figures. O.B performed the genomic sequencing. J.W and M.W.M designed the experiments and supervised the study. A.S.G and M.W.M wrote the manuscript with inputs from all co-authors.

## Acknowledgements

We thank Drs Elke Ströher and Owen Duncan (WA Proteomics Facility, Centre for Microscopy, Characterisation, and Analysis, UWA) for their assistance in proteomic turnover rate analysis.

## Parsed Citations

**Arlt, H., Steglich, G., Perryman, R., Guiard, B., Neupert, W., and Langer, T. (1998). The formation of respiratory chain complexes in mitochondria is under the proteolytic control of the m-AAA protease. EMBO J. 17: 4837–4847.**

Google Scholar: <u>Author Only Title Only Author and Title</u>

**Baradaran, R., Berrisford, J.M., Minhas, G.S., and Sazanov, L.A. (2013). Crystal structure of the entire respiratory complex i. Nature 494: 443–448.**

Google Scholar: <u>Author Only Title Only Author and Title</u>

**Bateman, A. et al. (2017). UniProt: The universal protein knowledgebase. Nucleic Acids Res. 45: D158–D169.**

Google Scholar: <u>Author Only Title Only Author and Title</u>

**Bindea, G., Mlecnik, B., Hackl, H., Charoentong, P., Tosolini, M., Kirilovsky, A., Fridman, W.H., Pagès, F., Trajanoski, Z., and Galon, J. (2009). ClueGO: A Cytoscape plug-in to decipher functionally grouped gene ontology and pathway annotation networks. Bioinformatics 25: 1091–1093.**

Google Scholar: <u>Author Only Title Only Author and Title</u>

**Branca, R.M.M., Orre, L.M., Johansson, H.J., Granholm, V., Huss, M., Pérez-Bercoff, A., Forshed, J., Käll, L., and Lehtiö, J. (2014). HiRIEF LC-MS enables deep proteome coverage and unbiased proteogenomics. Nat. Methods 11: 59–62.**

Google Scholar: <u>Author Only Title Only Author and Title</u>

**Carrie, C., Giraud, E., Duncan, O., Xu, L., Wang, Y., Huang, S., Clifton, R., Murcha, M., Filipovska, A., Rackham, O., Vrielink, A., and Whelan, J. (2010). Conserved and novel functions for Arabidopsis thaliana MIA40 in assembly of proteins in mitochondria and peroxisomes. J. Biol. Chem. 285: 36138–36148.**

Google Scholar: <u>Author Only Title Only Author and Title</u>

**Citovsky, V., Lee, L.Y., Vyas, S., Glick, E., Chen, M.H., Vainstein, A., Gafni, Y., Gelvin, S.B., and Tzfira, T. (2006). Subcellular**

**Localization of Interacting Proteins by Bimolecular Fluorescence Complementation in Planta. J. Mol. Biol. 362: 1120–1131.**

Google Scholar: <u>Author Only Title Only Author and Title</u>

**Duncan, O., Millar, A.H., and Taylor, N.L. (2017a). Isolation of Mitochondria, Their Sub-Organellar Compartments, and Membranes. In Isolation of Plant Organelles and Structures: Methods and Protocols, N.L. Taylor and A.H. Millar, eds (Springer New York: New York, NY), pp. 83–96.**

Google Scholar: <u>Author Only Title Only Author and Title</u>

**Duncan, O., Millar, A.H., and Taylor, N.L. (2017b). Isolation of Mitochondria, Their Sub-Organellar Compartments, and Membranes BT - Isolation of Plant Organelles and Structures: Methods and Protocols. In N.L. Taylor and A.H. Millar, eds (Springer New York: New York, NY), pp. 83–96.**

Google Scholar: <u>Author Only Title Only Author and Title</u>

**Dutilleul, C., Garmier, M., Noctor, G., Mathieu, C., Chétrit, P., Foyer, C.H., and De Paepe, R. (2003). Leaf mitochondria modulate whole cell redox homeostasis, set antioxidant capacity, and determine stress resistance through altered signaling and diurnal regulation. Plant Cell 15: 1212–1226.**

Google Scholar: <u>Author Only Title Only Author and Title</u>

**Eubel, H., Braun, H.P., and Millar, A.H. (2005). Blue-native PAGE in plants: A tool in analysis of protein-protein interactions. Plant Methods 1: 1–13.**

Google Scholar: <u>Author Only Title Only Author and Title</u>

**Fiedorczuk, K., Letts, J.A., Degliesposti, G., Kaszuba, K., Skehel, M., and Sazanov, L.A. (2016). Atomic structure of the entire mammalian mitochondrial complex i. Nature 538: 406–410.**

Google Scholar: <u>Author Only Title Only Author and Title</u>

**Fromm, S., Senkler, J., Eubel, H., Peterhänsel, C., and Braun, H.P. (2016). Life without complex I: Proteome analyses of an Arabidopsis mutant lacking the mitochondrial NADH dehydrogenase complex. J. Exp. Bot. 67: 3079–3093.**

Google Scholar: <u>Author Only Title Only Author and Title</u>

**Galemou Yoga, E., Haapanen, O., Wittig, I., Siegmund, K., Sharma, V., and Zickermann, V. (2019). Mutations in a conserved loop in the PSST subunit of respiratory complex I affect ubiquinone binding and dynamics. Biochim. Biophys. Acta - Bioenerg. 1860: 573– 581.**

Google Scholar: <u>Author Only Title Only Author and Title</u>

**Ghifari, A.S. and Murcha, M.W. (2022). Proteolytic regulation of mitochondrial oxidative phosphorylation components in plants. Biochem. Soc. Trans. 1: 1–14.**

Google Scholar: <u>Author Only Title Only Author and Title</u>

**Herman, C., Prakash, S., Lu, C.Z., Matouschek, A., and Gross, C.A. (2003). Lack of a robust unfoldase activity confers a unique level of substrate specificity to the universal AAA protease FtsH. Mol. Cell 11: 659–669.**

Google Scholar: <u>Author Only Title Only Author and Title</u>

**Hirst, J. and Roessler, M.M. (2016). Energy conversion, redox catalysis and generation of reactive oxygen species by respiratory complex i. Biochim. Biophys. Acta - Bioenerg. 1857: 872–883.**

Google Scholar: <u>Author Only Title Only Author and Title</u>

**Huala, E. et al. (2001). The Arabidopsis Information Resource (TAIR): a comprehensive database and web-based information retrieval, analysis, and visualization system for a model plant. Nucleic Acids Res. 29: 102–105.**

Google Scholar: <u>Author Only Title Only Author and Title</u>

**Huang, S., Lee, C.P., and Millar, A.H. (2015). Activity Assay for Plant Mitochondrial Enzymes. In Plant Mitochondria: Methods and Protocols, J. Whelan and M.W. Murcha, eds (Springer New York: New York, NY), pp. 139–149.**

Google Scholar: <u>Author Only Title Only Author and Title</u>

**Huang, S., Petereit, J., and Millar, A.H. (2020). Loss of conserved mitochondrial CLPP and its functions lead to different phenotypes in plants and other organisms. Plant Signal. Behav. 15.**

Google Scholar: <u>Author Only Title Only Author and Title</u>

**Ivanova, A., Ghifari, A.S., Berkowitz, O., Whelan, J., and Murcha, M.W. (2021). The mitochondrial AAA protease FTSH3 regulates Complex I abundance by promoting its disassembly. Plant Physiol. 186: 599–610.**

Google Scholar: <u>Author Only Title Only Author and Title</u>

**Ivanova, A., Gill-Hille, M., Huang, S., Branca, R., Kmiec, B., Teixeira, P.F., Lehtiö, J., Whelan, J., and Murcha, M.W. (2019). A Mitochondrial LYR Protein is Required for Complex I Assembly. Plant Physiol. 181: 1632–1650.**

Google Scholar: <u>Author Only Title Only Author and Title</u>

**Karimi, M., Inzé, D., and Depicker, A. (2002). GATEWAY vectors for Agrobacterium-mediated plant transformation. Trends Plant Sci. 7: 193–195.**

Google Scholar: <u>Author Only Title Only Author and Title</u>

**Kato, Y., Miura, E., Ido, K., Ifuku, K., and Sakamoto, W. (2009). The variegated mutants lacking chloroplastic FtsHs are defective in D1 degradation and accumulate reactive oxygen species. Plant Physiol. 151: 1790–1801.**

Google Scholar: <u>Author Only Title Only Author and Title</u>

**Katzen, F. (2007). Gateway^®^ recombinational cloning: A biological operating system. Expert Opin. Drug Discov. 2: 571–589.**

Google Scholar: <u>Author Only Title Only Author and Title</u>

**Kelley, L.A., Mezulis, S., Yates, C., Wass, M., and Sternberg, M. (2015). The Phyre2 web portal for protein modelling, prediction, and analysis. Nat. Protoc. 10: 845–858.**

Google Scholar: <u>Author Only Title Only Author and Title</u>

**Kihara, A., Akiyama, Y., and Ito, K. (1995). FtsH is required for proteolytic elimination of uncomplexed forms of SecY, an essential protein translocase subunit. Proc. Natl. Acad. Sci. U. S. A. 92: 4532–4536.**

Google Scholar: <u>Author Only Title Only Author and Title</u>

**Kihara, A., Akiyama, Y., and Koreaki, I. (1999). Dislocation of membrane proteins in FtsH-mediated proteolysis. EMBO J. 18: 2970– 2981.**

Google Scholar: <u>Author Only Title Only Author and Title</u>

**Klusch, N., Dreimann, M., Senkler, J., Rugen, N., Kühlbrandt, W., and Braun, H.P. (2022). Cryo-EM structure of the respiratory I + III2 supercomplex from Arabidopsis thaliana at 2 Å resolution. Nat. Plants.**

Google Scholar: <u>Author Only Title Only Author and Title</u>

**Klusch, N., Senkler, J., Yildiz, Ö., Kühlbrandt, W., and Braun, H.-P. (2021). A ferredoxin bridge connects the two arms of plant mitochondrial complex I. Plant Cell 33: 2072–2091.**

Google Scholar: <u>Author Only Title Only Author and Title</u>

**Kmiec, B., Branca, R.M.M., Berkowitz, O., Li, L., Wang, Y., Murcha, M.W., Whelan, J., Lehtiö, J., Glaser, E., and Teixeira, P.F. (2018). Accumulation of endogenous peptides triggers a pathogen stress response in Arabidopsis thaliana. Plant J. 96: 705–715.**

Google Scholar: <u>Author Only Title Only Author and Title</u>

**Kolata, P. and Efremov, R.G. (2021). Structure of Escherichia coli respiratory complex I reconstituted into lipid nanodiscs reveals an uncoupled conformation. Elife 10: 1–32.**

Google Scholar: <u>Author Only Title Only Author and Title</u>

**Kolodziejczak, M., Gibala, M., Urantowka, A., and Janska, H. (2007). The significance of Arabidopsis AAA proteases for activity and assembly/stability of mitochondrial OXPHOS complexes. Physiol. Plant. 129: 135–142.**

Google Scholar: <u>Author Only Title Only Author and Title</u>

**Kolodziejczak, M., Skibior-Blaszczyk, R., and Janska, H. (2018). m-AAA Complexes Are Not Crucial for the Survival of Arabidopsis under Optimal Growth Conditions Despite Their Importance for Mitochondrial Translation. Plant Cell Physiol. 59: 1006–1016.**

Google Scholar: <u>Author Only Title Only Author and Title</u>

**Koppen, M., Metodiev, M.D., Casari, G., Rugarli, E.I., and Langer, T. (2007). Variable and Tissue-Specific Subunit Composition of Mitochondrial m -AAA Protease Complexes Linked to Hereditary Spastic Paraplegia. Mol. Cell. Biol. 27: 758–767.**

Google Scholar: <u>Author Only Title Only Author and Title</u>

**Kühn, K., Obata, T., Feher, K., Bock, R., Fernie, A.R., and Meyer, E.H. (2015). Complete mitochondrial complex I deficiency induces an up-regulation of respiratory fluxes that is abolished by traces of functional complex I. Plant Physiol. 168: 1537–1549.**

Google Scholar: <u>Author Only Title Only Author and Title</u>

**Langmead, B. and Salzberg, S.L. (2012). Fast gapped-read alignment with Bowtie 2. Nat. Methods 9: 357–359.**

Google Scholar: <u>Author Only Title Only Author and Title</u>

**Li, L., Nelson, C., Fenske, R., Trösch, J., Pružinská, A., Millar, A.H., and Huang, S. (2017). Changes in specific protein degradation rates in Arabidopsis thaliana reveal multiple roles of Lon1 in mitochondrial protein homeostasis. Plant J. 89: 458–471.**

Google Scholar: <u>Author Only Title Only Author and Title</u>

**Ligas, J., Pineau, E., Bock, R., Huynen, M.A, and Meyer, E.H. (2019). The assembly pathway of complex I in Arabidopsis thaliana. Plant J. 97: 447–459.**

Google Scholar: <u>Author Only Title Only Author and Title</u>

**Maldonado, M., Fan, Z., Abe, K.M., and Letts, J.A. (2022). Plant-specific features of respiratory supercomplex I + III2 from Vigna radiata. Nat. Plants 2.**

Google Scholar: <u>Author Only Title Only Author and Title</u>

**Maldonado, M., Padavannil, A., Zhou, L., Guo, F., and Letts, J.A. (2020). Atomic structure of a mitochondrial complex I intermediate from vascular plants. Elife 9: e56664.**

Google Scholar: <u>Author Only Title Only Author and Title</u>

**Martin, A., Baker, T.A., and Sauer, R.T. (2008). Protein unfolding by a AAA+ protease is dependent on ATP-hydrolysis rates and substrate energy landscapes. Nat. Struct. Mol. Biol. 15: 139–145.**

Google Scholar: <u>Author Only Title Only Author and Title</u>

**Maziak, A., Heidorn-Czarna, M., Weremczuk, A., and Janska, H. (2021). FTSH4 and OMA1 mitochondrial proteases reduce moderate heat stress-induced protein aggregation. Plant Physiol. 187: 1-18.**

Google Scholar: <u>Author Only Title Only Author and Title</u>

**Meyer, E.H., Solheim, C., Tanz, S.K., Bonnard, G., and Millar, A.H. (2011). Insights into the composition and assembly of the membrane arm of plant complex I through analysis of subcomplexes in Arabidopsis mutant lines. J. Biol. Chem. 286: 26081–26092.**

Google Scholar: <u>Author Only Title Only Author and Title</u>

**Meyer, E.H., Tomaz, T., Carroll, A.J., Estavillo, G., Delannoy, E., Tanz, S.K., Small, I.D., Pogson, B.J., and Millar, A.H. (2009). Remodeled respiration in ndufs4 with Low phosphorylation efficiency suppresses Arabidopsis germination and growth and alters control of metabolism at night. Plant Physiol. 151: 603–619.**

Google Scholar: <u>Author Only Title Only Author and Title</u>

**Murcha, M.W., Lister, R., Ho, A.Y., and Whelan, J. (2003). Identification, expression, and import of components 17 and 23 of the inner mitochondrial membrane translocase from Arabidopsis. Plant Physiol 131: 1737–1747.**

Google Scholar: <u>Author Only Title Only Author and Title</u>

**Nakagawa, T., Kurose, T., Hino, T., Tanaka, K., Kawamukai, M., Niwa, Y., Toyooka, K., Matsuoka, K., Jinbo, T., and Kimura, T. (2007). Development of series of gateway binary vectors, pGWBs, for realizing efficient construction of fusion genes for plant transformation. J. Biosci. Bioeng. 104: 34–41.**

Google Scholar: <u>Author Only Title Only Author and Title</u>

**Nelson, C.J., Li, L., Jacoby, R.P., and Millar, A.H. (2013). Degradation rate of mitochondrial proteins in Arabidopsis thaliana cells. J. Proteome Res. 12: 3449–3459.**

Google Scholar: <u>Author Only Title Only Author and Title</u>

**Opalińska, M. and Jańska, H. (2018). AAA Proteases: Guardians of Mitochondrial Function and Homeostasis. Cells 7: 163.**

Google Scholar: <u>Author Only Title Only Author and Title</u>

**Parey, K., Haapanen, O., Sharma, V., Köfeler, H., Züllig, T., Prinz, S., Siegmund, K., Wittig, I., Mills, D.J., Vonck, J., Kühlbrandt, W., and Zickermann, V. (2019). High-resolution cryo-EM structures of respiratory complex I: Mechanism, assembly, and disease. Sci. Adv. 5: 1–11.**

Google Scholar: <u>Author Only Title Only Author and Title</u>

**Perales, M., Eubel, H., Heinemeyer, J., Colaneri, A., Zabaleta, E., and Braun, H.P. (2005). Disruption of a nuclear gene encoding a mitochondrial gamma carbonic anhydrase reduces complex I and supercomplex I+III2 levels and alters mitochondrial physiology in Arabidopsis. J. Mol. Biol. 350: 263–277.**

Google Scholar: <u>Author Only Title Only Author and Title</u>

**Petereit, J. et al. (2020). Mitochondrial CLPP2 Assists Coordination and Homeostasis of Respiratory Complexes. Plant Physiol. 184: 148–164.**

Google Scholar: <u>Author Only Title Only Author and Title</u>

**Peters, K., Niessen, M., Peterhansel, C., Spath, B., Holzle, A., Binder, S., Marchfelder, A., and Braun, H.P. (2012). Complex I-complex II ratio strongly differs in various organs of Arabidopsis thaliana. Plant Mol Biol 79: 273–284.**

Google Scholar: <u>Author Only Title Only Author and Title</u>

**Pettersen, E.F., Goddard, T.D., Huang, C.C., Meng, E.C., Couch, G.S., Croll, T.I., Morris, J.H., and Ferrin, T.E. (2021). UCSF ChimeraX: Structure visualization for researchers, educators, and developers. Protein Sci. 30: 70–82.**

Google Scholar: <u>Author Only Title Only Author and Title</u>

**Piechota, J., Kolodziejczak, M., Juszczak, I., Sakamoto, W., and Janska, H. (2010). Identification and Characterization of High Molecular Weight Complexes Formed by Matrix AAA Proteases and Prohibitins in Mitochondria of Arabidopsis thaliana. J. Biol. Chem. 285: 12512–12521.**

Google Scholar: <u>Author Only Title Only Author and Title</u>

**Puchades, C., Ding, B., Song, A., Wiseman, R.L., Lander, G.C., and Glynn, S.E. (2019). Unique Structural Features of the Mitochondrial AAA+ Protease AFG3L2 Reveal the Molecular Basis for Activity in Health and Disease. Mol. Cell 75: 1073–1085.e6.**

Google Scholar: <u>Author Only Title Only Author and Title</u>

**Puchades, C., Rampello, A.J., Shin, M., Giuliano, C.J., Wiseman, R.L., Glynn, S.E., and Lander, G.C. (2017). Structure of the mitochondrial inner membrane AAA+ protease YME1 gives insight into substrate processing. Science (80-.). 358: eaao0464.**

Google Scholar: <u>Author Only Title Only Author and Title</u>

**Schertl, P. and Braun, H.-P. (2015). Activity Measurements of Mitochondrial Enzymes in Native Gels. In Plant Mitochondria: Methods and Protocols, J. Whelan and M.W. Murcha, eds (Springer New York: New York, NY), pp. 131–138.**

Google Scholar: <u>Author Only Title Only Author and Title</u>

**Schimmeyer, J., Bock, R., and Meyer, E.H. (2016). l-Galactono-1,4-lactone dehydrogenase is an assembly factor of the membrane arm of mitochondrial complex I in Arabidopsis. Plant Mol. Biol. 90: 117–126.**

Google Scholar: <u>Author Only Title Only Author and Title</u>

**Schneider, C.A, Rasband, W.S., and Eliceiri, K.W. (2012). NIH Image to ImageJ: 25 years of image analysis. Nat. Methods 9: 671– 675.**

Google Scholar: <u>Author Only Title Only Author and Title</u>

**Shannon, P., Markiel, A., Ozier, O., Baliga, N.S., Wang, J.T., Ramage, D., Amin, N., Schwikowski, B., and Ideker, T. (2003). Cytoscape: A Software Environment for Integrated Models. Genome Res. 13: 2498–2504.**

Google Scholar: <u>Author Only Title Only Author and Title</u>

**Shi, H., Rampello, A.J., and Glynn, S.E. (2016). Engineered AAA+ proteases reveal principles of proteolysis at the mitochondrial inner membrane. Nat. Commun. 7: 13301.**

Google Scholar: <u>Author Only Title Only Author and Title</u>

**Shin, M., Puchades, C., Asmita, A., Puri, N., Adjei, E., Wiseman, R.L., Karzai, A.W., and Lander, G.C. (2020). Structural basis for distinct operational modes and protease activation in AAA+ protease Lon. Sci. Adv. 6: 1–16.**

Google Scholar: <u>Author Only Title Only Author and Title</u>

**Shin, M., Watson, E.R., Song, A.S., Mindrebo, J.T., Novick, S.J., Griffin, P.R., Wiseman, R.L., and Lander, G.C. (2021). Structures of the human LONP1 protease reveal regulatory steps involved in protease activation. Nat. Commun. 12.**

Google Scholar: <u>Author Only Title Only Author and Title</u>

**Sievers, F., Wilm, A., Dineen, D., Gibson, T.J., Karplus, K., Li, W., Lopez, R., McWilliam, H., Remmert, M., Soding, J., Thompson, J.D., and Higgins, D.G. (2011). Fast, scalable generation of high-quality protein multiple sequence alignments using Clustal Omega. Mol. Syst. Biol. 7: 539.**

Google Scholar: <u>Author Only Title Only Author and Title</u>

**Silva, P., Thompson, E., Bailey, S., Kruse, O., Mullineaux, C.W., Robinson, C., Mann, N.H., and Nixon, P.J. (2003). FtsH Is Involved in the Early Stages of Repair of Photosystem II in Synechocystis sp PCC 6803. Plant Cell 15: 2152–2164.**

Google Scholar: <u>Author Only Title Only Author and Title</u>

**Soufari, H., Parrot, C., Kuhn, L., Waltz, F., and Hashem, Y. (2020). Specific features and assembly of the plant mitochondrial complex I revealed by cryo-EM. Nat. Commun. 11: 5195.**

Google Scholar: <u>Author Only Title Only Author and Title</u>

**Stellberger, T., Häuser, R., Baiker, A., Pothineni, V.R., Haas, J., and Uetz, P. (2010). Improving the yeast two-hybrid system with permutated fusions proteins: the Varicella Zoster Virus interactome. Proteome Sci. 8: 8.**

Google Scholar: <u>Author Only Title Only Author and Title</u>

**Subrahmanian, N., Remacle, C., and Hamel, P.P. (2016). Plant mitochondrial Complex I composition and assembly: A review. Biochim. Biophys. Acta - Bioenerg. 1857: 1001–1014.**

Google Scholar: <u>Author Only Title Only Author and Title</u>

**Szczepanowska, K. et al. (2020). A salvage pathway maintains highly functional respiratory complex I. Nat. Commun. 11: 1643.**

Google Scholar: <u>Author Only Title Only Author and Title</u>

**Urantowka, A., Knorpp, C., Olczak, T., Kolodziejczak, M., and Janska, H. (2005). Plant mitochondria contain at least two i-AAA-like complexes. Plant Mol. Biol. 59: 239–252.**

Google Scholar: <u>Author Only Title Only Author and Title</u>

**Vierstra, R.D. (2009). The ubiquitin-26S proteasome system at the nexus of plant biology. Nat. Rev. Mol. Cell Biol. 10: 385–397.**

Google Scholar: <u>Author Only Title Only Author and Title</u>

**Wing, Y., Carrie, C., Giraud, E., Elhafez, D., Narsai, R., Duncan, O., Whelan, J., and Murcha, M.W. (2012). Dual Location of the Mitochondrial Preprotein Transporters B14.7 and Tim23-2 in Complex I and the TIM17:23 Complex in Arabidopsis Links Mitochondrial Activity and Biogenesis. Plant Cell 24: 2675–2695.**

Google Scholar: <u>Author Only Title Only Author and Title</u>

**Weigel, D. and Glazebrook, J. (2006). EMS Mutagenesis of Arabidopsis Seed. Cold Spring Harb. Protoc. 2006: pdb.prot4621.**

Google Scholar: <u>Author Only Title Only Author and Title</u>

**van Wijk, K.J. (2015). Protein Maturation and Proteolysis in Plant Plastids, Mitochondria, and Peroxisomes. Annu. Rev. Plant Biol. 66: 75–111.**

Google Scholar: <u>Author Only Title Only Author and Title</u>

**Wu, F.H., Shen, S.C., Lee, L.Y., Lee, S.H., Chan, M.T., and Lin, C.S. (2009). Tape-arabidopsis sandwich - A simpler arabidopsis protoplast isolation method. Plant Methods 5: 1–10.**

Google Scholar: <u>Author Only Title Only Author and Title</u>

**Yoo, S.D., Cho, Y.H., and Sheen, J. (2007). Arabidopsis mesophyll protoplasts: A versatile cell system for transient gene expression analysis. Nat. Protoc. 2: 1565–1572.**

Google Scholar: <u>Author Only Title Only Author and Title</u>

## References

Arlt, H., Steglich, G., Perryman, R., Guiard, B., Neupert, W., and Langer, T. (1998). The formation of respiratory chain complexes in mitochondria is under the proteolytic control of the m-AAA protease. EMBO J. 17: 4837–4847.

Baradaran, R., Berrisford, J.M., Minhas, G.S., and Sazanov, L.A. (2013). Crystal structure of the entire respiratory complex i. Nature 494: 443–448.

Bateman, A. et al. (2017). UniProt: The universal protein knowledgebase. Nucleic Acids Res. 45: D158–D169.

Bindea, G., Mlecnik, B., Hackl, H., Charoentong, P., Tosolini, M., Kirilovsky, A., Fridman, W.H., Pagès, F., Trajanoski, Z., and Galon, J. (2009). ClueGO: A Cytoscape plug-in to decipher functionally grouped gene ontology and pathway annotation networks. Bioinformatics 25: 1091–1093.

Branca, R.M.M., Orre, L.M., Johansson, H.J., Granholm, V., Huss, M., Pérez-Bercoff, A., Forshed, J., Käll, L., and Lehtiö, J. (2014). HiRIEF LC-MS enables deep proteome coverage and unbiased proteogenomics. Nat. Methods 11: 59–62.

Carrie, C., Giraud, E., Duncan, O., Xu, L., Wang, Y., Huang, S., Clifton, R., Murcha, M., Filipovska, A., Rackham, O., Vrielink, A., and Whelan, J. (2010). Conserved and novel functions for Arabidopsis thaliana MIA40 in assembly of proteins in mitochondria and peroxisomes. J. Biol. Chem. 285: 36138–36148.

Citovsky, V., Lee, L.Y., Vyas, S., Glick, E., Chen, M.H., Vainstein, A., Gafni, Y., Gelvin, S.B., and Tzfira, T. (2006). Subcellular Localization of Interacting Proteins by Bimolecular Fluorescence Complementation in Planta. J. Mol. Biol. 362: 1120–1131.

Duncan, O., Millar, A.H., and Taylor, N.L. (2017a). Isolation of Mitochondria, Their Sub-Organellar Compartments, and Membranes. In Isolation of Plant Organelles and Structures: Methods and Protocols, N.L. Taylor and A.H. Millar, eds (Springer New York: New York, NY), pp. 83–96.

Duncan, O., Millar, A.H., and Taylor, N.L. (2017b). Isolation of Mitochondria, Their Sub-Organellar Compartments, and Membranes BT - Isolation of Plant Organelles and Structures: Methods and Protocols. In N.L. Taylor and A.H. Millar, eds (Springer New York: New York, NY), pp. 83–96.

Dutilleul, C., Garmier, M., Noctor, G., Mathieu, C., Chétrit, P., Foyer, C.H., and De Paepe, R. (2003). Leaf mitochondria modulate whole cell redox homeostasis, set antioxidant capacity, and determine stress resistance through altered signaling and diurnal regulation. Plant Cell 15: 1212–1226.

Eubel, H., Braun, H.P., and Millar, A.H. (2005). Blue-native PAGE in plants: A tool in analysis of protein-protein interactions. Plant Methods 1: 1–13.

Fiedorczuk, K., Letts, J.A., Degliesposti, G., Kaszuba, K., Skehel, M., and Sazanov, L.A. (2016). Atomic structure of the entire mammalian mitochondrial complex i. Nature 538: 406–410.

Fromm, S., Senkler, J., Eubel, H., Peterhänsel, C., and Braun, H.P. (2016). Life without complex I: Proteome analyses of an Arabidopsis mutant lacking the mitochondrial NADH dehydrogenase complex. J. Exp. Bot. 67: 3079–3093.

Galemou Yoga, E., Haapanen, O., Wittig, I., Siegmund, K., Sharma, V., and Zickermann, V. (2019). Mutations in a conserved loop in the PSST subunit of respiratory complex I affect ubiquinone binding and dynamics. Biochim. Biophys. Acta - Bioenerg. 1860: 573–581.

Ghifari, A.S. and Murcha, M.W. (2022). Proteolytic regulation of mitochondrial oxidative phosphorylation components in plants. Biochem. Soc. Trans. 1: 1–14.

Herman, C., Prakash, S., Lu, C.Z., Matouschek, A., and Gross, C.A. (2003). Lack of a robust unfoldase activity confers a unique level of substrate specificity to the universal AAA protease FtsH. Mol. Cell 11: 659–669.

Hirst, J. and Roessler, M.M. (2016). Energy conversion, redox catalysis and generation of reactive oxygen species by respiratory complex i. Biochim. Biophys. Acta - Bioenerg. 1857: 872–883.

Huala, E. et al. (2001). The Arabidopsis Information Resource (TAIR): a comprehensive database and web-based information retrieval, analysis, and visualization system for a model plant. Nucleic Acids Res. 29: 102–105.

Huang, S., Lee, C.P., and Millar, A.H. (2015). Activity Assay for Plant Mitochondrial Enzymes. In Plant Mitochondria: Methods and Protocols, J. Whelan and M.W. Murcha, eds (Springer New York: New York, NY), pp. 139–149.

Huang, S., Petereit, J., and Millar, A.H. (2020). Loss of conserved mitochondrial CLPP and its functions lead to different phenotypes in plants and other organisms. Plant Signal. Behav. 15.

Ivanova, A., Ghifari, A.S., Berkowitz, O., Whelan, J., and Murcha, M.W. (2021). The mitochondrial AAA protease FTSH3 regulates Complex I abundance by promoting its disassembly. Plant Physiol. 186: 599–610.

Ivanova, A., Gill-Hille, M., Huang, S., Branca, R., Kmiec, B., Teixeira, P.F., Lehtiö, J., Whelan, J., and Murcha, M.W. (2019). A Mitochondrial LYR Protein is Required for Complex I Assembly. Plant Physiol. 181: 1632–1650.

Karimi, M., Inzé, D., and Depicker, A. (2002). GATEWAY vectors for Agrobacterium-mediated plant transformation. Trends Plant Sci. 7: 193–195.

Kato, Y., Miura, E., Ido, K., Ifuku, K., and Sakamoto, W. (2009). The variegated mutants lacking chloroplastic FtsHs are defective in D1 degradation and accumulate reactive oxygen species. Plant Physiol. 151: 1790–1801.

Katzen, F. (2007). Gateway^®^ recombinational cloning: A biological operating system. Expert Opin. Drug Discov. 2: 571–589.

Kelley, L.A., Mezulis, S., Yates, C., Wass, M., and Sternberg, M. (2015). The Phyre2 web portal for protein modelling, prediction, and analysis. Nat. Protoc. 10: 845–858.

Kihara, A., Akiyama, Y., and Ito, K. (1995). FtsH is required for proteolytic elimination of uncomplexed forms of SecY, an essential protein translocase subunit. Proc. Natl. Acad. Sci. U. S. A. 92: 4532–4536.

Kihara, A., Akiyama, Y., and Koreaki, I. (1999). Dislocation of membrane proteins in FtsH-mediated proteolysis. EMBO J. 18: 2970–2981.

Klusch, N., Dreimann, M., Senkler, J., Rugen, N., Kühlbrandt, W., and Braun, H.P. (2022). Cryo-EM structure of the respiratory I + III2 supercomplex from Arabidopsis thaliana at 2 Å resolution. Nat. Plants.

Klusch, N., Senkler, J., Yildiz, Ö., Kühlbrandt, W., and Braun, H.-P. (2021). A ferredoxin bridge connects the two arms of plant mitochondrial complex I. Plant Cell 33: 2072–2091.

Kmiec, B., Branca, R.M.M., Berkowitz, O., Li, L., Wang, Y., Murcha, M.W., Whelan, J., Lehtiö, J., Glaser, E., and Teixeira, P.F. (2018). Accumulation of endogenous peptides triggers a pathogen stress response in *Arabidopsis thaliana*. Plant J. 96: 705–715.

Kolata, P. and Efremov, R.G. (2021). Structure of Escherichia coli respiratory complex I reconstituted into lipid nanodiscs reveals an uncoupled conformation. Elife 10: 1–32.

Kolodziejczak, M., Gibala, M., Urantowka, A., and Janska, H. (2007). The significance of Arabidopsis AAA proteases for activity and assembly/stability of mitochondrial OXPHOS complexes. Physiol. Plant. 129: 135–142.

Kolodziejczak, M., Skibior-Blaszczyk, R., and Janska, H. (2018). m-AAA Complexes Are Not Crucial for the Survival of Arabidopsis under Optimal Growth Conditions Despite Their Importance for Mitochondrial Translation. Plant Cell Physiol. 59: 1006–1016.

Koppen, M., Metodiev, M.D., Casari, G., Rugarli, E.I., and Langer, T. (2007). Variable and Tissue-Specific Subunit Composition of Mitochondrial m -AAA Protease Complexes Linked to Hereditary Spastic Paraplegia. Mol. Cell. Biol. 27: 758–767.

Kühn, K., Obata, T., Feher, K., Bock, R., Fernie, A.R., and Meyer, E.H. (2015). Complete mitochondrial complex I deficiency induces an up-regulation of respiratory fluxes that is abolished by traces of functional complex I. Plant Physiol. 168: 1537–1549.

Langmead, B. and Salzberg, S.L. (2012). Fast gapped-read alignment with Bowtie 2. Nat. Methods 9: 357–359.

Li, L., Nelson, C., Fenske, R., Trösch, J., Pružinská, A., Millar, A.H., and Huang, S. (2017). Changes in specific protein degradation rates in Arabidopsis thaliana reveal multiple roles of Lon1 in mitochondrial protein homeostasis. Plant J. 89: 458–471.

Ligas, J., Pineau, E., Bock, R., Huynen, M.A., and Meyer, E.H. (2019). The assembly pathway of complex I in Arabidopsis thaliana. Plant J. 97: 447–459.

Maldonado, M., Fan, Z., Abe, K.M., and Letts, J.A. (2022). Plant-specific features of respiratory supercomplex I + III2 from Vigna radiata. Nat. Plants 2.

Maldonado, M., Padavannil, A., Zhou, L., Guo, F., and Letts, J.A. (2020). Atomic structure of a mitochondrial complex I intermediate from vascular plants. Elife 9: e56664.

Martin, A., Baker, T.A., and Sauer, R.T. (2008). Protein unfolding by a AAA+ protease is dependent on ATP-hydrolysis rates and substrate energy landscapes. Nat. Struct. Mol. Biol. 15: 139–145.

Maziak, A., Heidorn-Czarna, M., Weremczuk, A., and Janska, H. (2021). FTSH4 and OMA1 mitochondrial proteases reduce moderate heat stress-induced protein aggregation. Plant Physiol. 187: 1–18.

Meyer, E.H., Solheim, C., Tanz, S.K., Bonnard, G., and Millar, A.H. (2011). Insights into the composition and assembly of the membrane arm of plant complex I through analysis of subcomplexes in Arabidopsis mutant lines. J. Biol. Chem. 286: 26081–26092.

Meyer, E.H., Tomaz, T., Carroll, A.J., Estavillo, G., Delannoy, E., Tanz, S.K., Small, I.D., Pogson, B.J., and Millar, A.H. (2009). Remodeled respiration in ndufs4 with Low phosphorylation efficiency suppresses Arabidopsis germination and growth and alters control of metabolism at night. Plant Physiol. 151: 603–619.

Murcha, M.W., Lister, R., Ho, A.Y., and Whelan, J. (2003). Identification, expression, and import of components 17 and 23 of the inner mitochondrial membrane translocase from Arabidopsis. Plant Physiol 131: 1737–1747.

Nakagawa, T., Kurose, T., Hino, T., Tanaka, K., Kawamukai, M., Niwa, Y., Toyooka, K., Matsuoka, K., Jinbo, T., and Kimura, T. (2007). Development of series of gateway binary vectors, pGWBs, for realizing efficient construction of fusion genes for plant transformation. J. Biosci. Bioeng. 104: 34–41.

Nelson, C.J., Li, L., Jacoby, R.P., and Millar, A.H. (2013). Degradation rate of mitochondrial proteins in Arabidopsis thaliana cells. J. Proteome Res. 12: 3449–3459.

Opalińska, M. and Jańska, H. (2018). AAA Proteases: Guardians of Mitochondrial Function and Homeostasis. Cells 7: 163.

Parey, K., Haapanen, O., Sharma, V., Köfeler, H., Züllig, T., Prinz, S., Siegmund, K., Wittig, I., Mills, D.J., Vonck, J., Kühlbrandt, W., and Zickermann, V. (2019). High-resolution cryo-EM structures of respiratory complex I: Mechanism, assembly, and disease. Sci. Adv. 5: 1–11.

Perales, M., Eubel, H., Heinemeyer, J., Colaneri, A., Zabaleta, E., and Braun, H.P. (2005). Disruption of a nuclear gene encoding a mitochondrial gamma carbonic anhydrase reduces complex I and supercomplex I+III2 levels and alters mitochondrial physiology in Arabidopsis. J. Mol. Biol. 350: 263–277.

Petereit, J. et al. (2020). Mitochondrial CLPP2 Assists Coordination and Homeostasis of Respiratory Complexes. Plant Physiol. 184: 148–164.

Peters, K., Niessen, M., Peterhansel, C., Spath, B., Holzle, A., Binder, S., Marchfelder, A., and Braun, H.P. (2012). Complex I-complex II ratio strongly differs in various organs of Arabidopsis thaliana. Plant Mol Biol 79: 273–284.

Pettersen, E.F., Goddard, T.D., Huang, C.C., Meng, E.C., Couch, G.S., Croll, T.I., Morris, J.H., and Ferrin, T.E. (2021). UCSF ChimeraX: Structure visualization for researchers, educators, and developers. Protein Sci. 30: 70–82.

Piechota, J., Kolodziejczak, M., Juszczak, I., Sakamoto, W., and Janska, H. (2010). Identification and Characterization of High Molecular Weight Complexes Formed by Matrix AAA Proteases and Prohibitins in Mitochondria of Arabidopsis thaliana. J. Biol. Chem. 285: 12512–12521.

Puchades, C., Ding, B., Song, A., Wiseman, R.L., Lander, G.C., and Glynn, S.E. (2019). Unique Structural Features of the Mitochondrial AAA+ Protease AFG3L2 Reveal the Molecular Basis for Activity in Health and Disease. Mol. Cell 75: 1073–1085.e6.

Puchades, C., Rampello, A.J., Shin, M., Giuliano, C.J., Wiseman, R.L., Glynn, S.E., and Lander, G.C. (2017). Structure of the mitochondrial inner membrane AAA+ protease YME1 gives insight into substrate processing. Science (80-.). 358: eaao0464.

Schertl, P. and Braun, H.-P. (2015). Activity Measurements of Mitochondrial Enzymes in Native Gels. In Plant Mitochondria: Methods and Protocols, J. Whelan and M.W. Murcha, eds (Springer New York: New York, NY), pp. 131–138.

Schimmeyer, J., Bock, R., and Meyer, E.H. (2016). l-Galactono-1,4-lactone dehydrogenase is an assembly factor of the membrane arm of mitochondrial complex I in Arabidopsis. Plant Mol. Biol. 90: 117–126.

Schneider, C.A., Rasband, W.S., and Eliceiri, K.W. (2012). NIH Image to ImageJ: 25 years of image analysis. Nat. Methods 9: 671–675.

Shannon, P., Markiel, A., Ozier, O., Baliga, N.S., Wang, J.T., Ramage, D., Amin, N., Schwikowski, B., and Ideker, T. (2003). Cytoscape: A Software Environment for Integrated Models. Genome Res. 13: 2498–2504.

Shi, H., Rampello, A.J., and Glynn, S.E. (2016). Engineered AAA+ proteases reveal principles of proteolysis at the mitochondrial inner membrane. Nat. Commun. 7: 13301.

Shin, M., Puchades, C., Asmita, A., Puri, N., Adjei, E., Wiseman, R.L., Karzai, A.W., and Lander, G.C. (2020). Structural basis for distinct operational modes and protease activation in AAA+ protease Lon. Sci. Adv. 6: 1–16.

Shin, M., Watson, E.R., Song, A.S., Mindrebo, J.T., Novick, S.J., Griffin, P.R., Wiseman, R.L., and Lander, G.C. (2021). Structures of the human LONP1 protease reveal regulatory steps involved in protease activation. Nat. Commun. 12.

Sievers, F., Wilm, A., Dineen, D., Gibson, T.J., Karplus, K., Li, W., Lopez, R., McWilliam, H., Remmert, M., Soding, J., Thompson, J.D., and Higgins, D.G. (2011). Fast, scalable generation of high-quality protein multiple sequence alignments using Clustal Omega. Mol. Syst. Biol. 7: 539.

Silva, P., Thompson, E., Bailey, S., Kruse, O., Mullineaux, C.W., Robinson, C., Mann, N.H., and Nixon, P.J. (2003). FtsH Is Involved in the Early Stages of Repair of Photosystem II in Synechocystis sp PCC 6803. Plant Cell 15: 2152–2164.

Soufari, H., Parrot, C., Kuhn, L., Waltz, F., and Hashem, Y. (2020). Specific features and assembly of the plant mitochondrial complex I revealed by cryo-EM. Nat. Commun. 11: 5195.

Stellberger, T., Häuser, R., Baiker, A., Pothineni, V.R., Haas, J., and Uetz, P. (2010). Improving the yeast two-hybrid system with permutated fusions proteins: the Varicella Zoster Virus interactome. Proteome Sci. 8: 8.

Subrahmanian, N., Remacle, C., and Hamel, P.P. (2016). Plant mitochondrial Complex I composition and assembly: A review. Biochim. Biophys. Acta - Bioenerg. 1857: 1001–1014.

Szczepanowska, K. et al. (2020). A salvage pathway maintains highly functional respiratory complex I. Nat. Commun. 11: 1643.

Urantowka, A., Knorpp, C., Olczak, T., Kolodziejczak, M., and Janska, H. (2005). Plant mitochondria contain at least two i-AAA-like complexes. Plant Mol. Biol. 59: 239–252.

Vierstra, R.D. (2009). The ubiquitin-26S proteasome system at the nexus of plant biology. Nat. Rev. Mol. Cell Biol. 10: 385–397.

Wang, Y., Carrie, C., Giraud, E., Elhafez, D., Narsai, R., Duncan, O., Whelan, J., and Murcha, M.W. (2012). Dual Location of the Mitochondrial Preprotein Transporters B14.7 and Tim23-2 in Complex I and the TIM17:23 Complex in Arabidopsis Links Mitochondrial Activity and Biogenesis. Plant Cell 24: 2675–2695.

Weigel, D. and Glazebrook, J. (2006). EMS Mutagenesis of Arabidopsis Seed. Cold Spring Harb. Protoc. 2006: pdb.prot4621.

van Wijk, K.J. (2015). Protein Maturation and Proteolysis in Plant Plastids, Mitochondria, and Peroxisomes. Annu. Rev. Plant Biol. 66: 75–111.

Wu, F.H., Shen, S.C., Lee, L.Y., Lee, S.H., Chan, M.T., and Lin, C.S. (2009). Tape-arabidopsis sandwich - A simpler arabidopsis protoplast isolation method. Plant Methods 5: 1–10.

Yoo, S.D., Cho, Y.H., and Sheen, J. (2007). Arabidopsis mesophyll protoplasts: A versatile cell system for transient gene expression analysis. Nat. Protoc. 2: 1565–1572.

